# Genetic elements and defense systems drive diversification and evolution in Asgard archaea

**DOI:** 10.1101/2024.03.22.586370

**Authors:** Luis E. Valentin-Alvarado, Ling-Dong Shi, Kathryn E. Appler, Alexander Crits-Christoph, Michael Cui, Valerie De Anda, Pedro Leão, Benjamin A. Adler, Richard J. Roberts, Rohan Sachdeva, Brett J. Baker, David F. Savage, Jillian F. Banfield

## Abstract

Asgard Archaea are of great interest as the progenitors of Eukaryotes, but little is known about the mobile genetic elements (MGEs) that may shape their ongoing evolution. Here, we describe MGEs that replicate in Atabeyarchaeia, wetland Asgard archaea phylum represented by two complete genomes. We used soil depth-resolved population metagenomic datasets to track 18 MGEs for which genome structures were defined and precise chromosome integration sites could be identified for confident host linkage. Additionally, we identified a complete 20.67 kilobase pair (kbp) circular plasmid (the first reported for Asgard archaea) and two groups of viruses linked to Atabeyarchaeia, via CRISPR spacer targeting. Closely related 40 kbp viruses possess a hypervariable genomic region encoding combinations of specific genes for small cysteine-rich proteins structurally similar to restriction-homing endonucleases. One 10.9 kbp circularizable plasmid-like MGE integrates genomically into an Atabeyarchaeia chromosome and has a 2.5 kbp circularizable element integrated within it. The 10.9 kbp MGE encodes a highly expressed methylase with a sequence specificity matching an active methylation motif identified by PacBio sequencing. Restriction-modification of Atabeyarchaeia differs from that of another coexisting Asgard archaea Freyarchaeia which has few identified MGEs but possesses diverse defense mechanisms, including DISARM and Hachiman not found in Atabeyarchaeia. Overall, defense systems and methylation mechanisms of Asgard archaea likely modulate their interactions with MGEs, and integration/excision and copy number variation of MGEs in turn enable host genetic versatility.

## Introduction

Asgard archaea, including Loki-, Hermod-, Thor-, Odin-, Baldr-, Freya-, Sif-, Heimdall-, Atabey-, and Wukongarchaeia, bridge our understanding of the evolution of eukaryotes and prokaryotes. Their genomic features, particularly the presence of eukaryotic signature proteins (ESPs), provide insights into the steps leading to eukaryotic cellular complexity. Recent phylogenetic analyses place eukaryotes within Asgard archaea(Spang et al., 2015; Zaremba-Niedzwiedzka et al., 2017), most closely related to Hodarchaeales(Eme et al., 2023). Despite intense interest in their functionality and evolutionary relationships, little has been reported regarding Asgard mobile genetic elements (MGEs) that may shape their population diversity, contribute to genome divergence and facilitate cross-domain horizontal gene transfer(Ghaly et al., 2022). Recent studies identified viruses of Loki-, Odin-, and Thor and Heimdallarchaeia (Medvedeva et al., 2022; Rambo et al., 2022; Tamarit et al., 2022; Wu et al., 2022), as well as putative transposons carrying cargo genes that replicate within Heimdallarchaeia(Wu et al., 2022), primarily based on CRISPR targeting. To our knowledge, no plasmids or plasmid-like elements have been described for Asgard archaea.

Recently, we reported two complete genomes for Atabeyarchaeia, a new group of Asgard archaea, and the first complete genome for Freyarchaeia(Valentin-Alvarado et al., 2023). Here, we track subtle strain variation of integrated MGEs over a soil depth profile by aligning the Illumina reads from a series of metagenomic samples from vernal pool wetland soil. We uncovered a suite of MGEs of Atabeyarchaeia that coexist in integrated and circular forms in metagenomic samples. Importantly, this approach establishes the host, and exactly defines MGE insertion sites and MGE lengths. Thus, we expand the repertoire of MGEs of Asgard archaea, explore their gene inventories, and shed light on MGE integration and excision events in natural populations of Atabeyarchaeia. Based on CRISPR targeting, we also genomically define groups of circular viruses and unclassified MGEs, some of which feature hypervariable regions enriched in cysteine-rich proteins predicted to be restriction endonucleases. We also describe genomically-encoded defense systems of both Atabeyarchaeia and Freyarchaeia that we confirm to be actively expressed with metatranscriptomic data. Leveraging the ability of PacBio SMRT sequencing data to identify DNA methylation sites, we report methylation patterns that distinguish these archaea, as well as a transcriptionally active MGE-encoded methylase that may enable the MGE to avoid restriction-based defense systems and compete with other MGEs. Overall, this study leverages complete metagenome-based genomes, read-based population analyses, metatranscriptomics and long read-based epigenetic analyses to provide a mosaic set of Asgard MGEs with likely relevance to Asgard archaeal evolution.

## Results

### Complete Atabeyarchaeia genomes contain integrated genetic mobile elements

Short and long-read metagenomic and metatranscriptomic datasets were generated from wetland soil sampled at a single local site (**Methods**). We mapped reads from 28 samples collected from soil depths of 60 to 175 cm to the Atabeyarchaeia-1, Atabeyarchaeia-2 and Freyarchaeia genomes previously assembled from this environment, and used mapped read details to uncover evidence for integrated, excised and coexisting circularized MGEs (**Figure 1**). Absence of MGEs in some cells can lead to lower coverage than average coverage over the integrated region, whereas higher read depth of coverage is likely due to coexisting extrachromosomal versions of the MGE(Kieft & Anantharaman, 2022). By manual inspection of sequencing depth and read alignment discrepancies, we identified 14 chromosomally integrated genetic elements in the Atabeyarchaeia-1 (Atabeya-1) genome and 4 in the Atabeyarchaeia-2 (Atabeya-2) genome, ranging from 1.3 to 40 kbp in length. No integrated elements were identified in the Freyarchaeia genomes using this approach (**Figure 2**).

**Figure 1:**
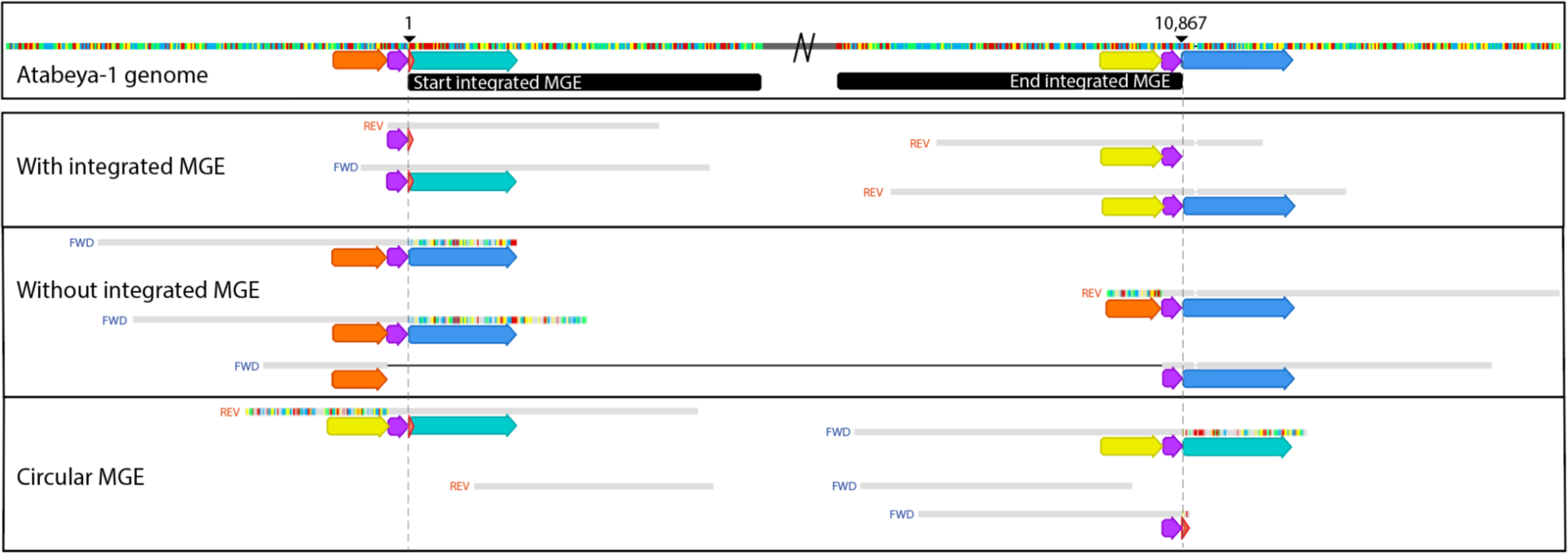
Read mapping to the reference genome provides evidence for integration and excision, illustrated for the case of one MGE. The central region of the integrated sequence of Yucahu-i (between black bars) has been deleted to focus on details of reads mapped to the start and end of the region. Read sequences that match the genome sequences are shown as gray bars; small vertical colored bars adjacent to read portions in agreement with the reference indicate bases that disagree with the reference. Arrows with the same color have the same nucleotide sequence. In the panel demonstrating that some cells that lack the integrated MGE, one read has been split (the black line links the two parts of a single read) to illustrate agreement with the flanking sequence at both ends of the integrated region.

**Figure 2.**
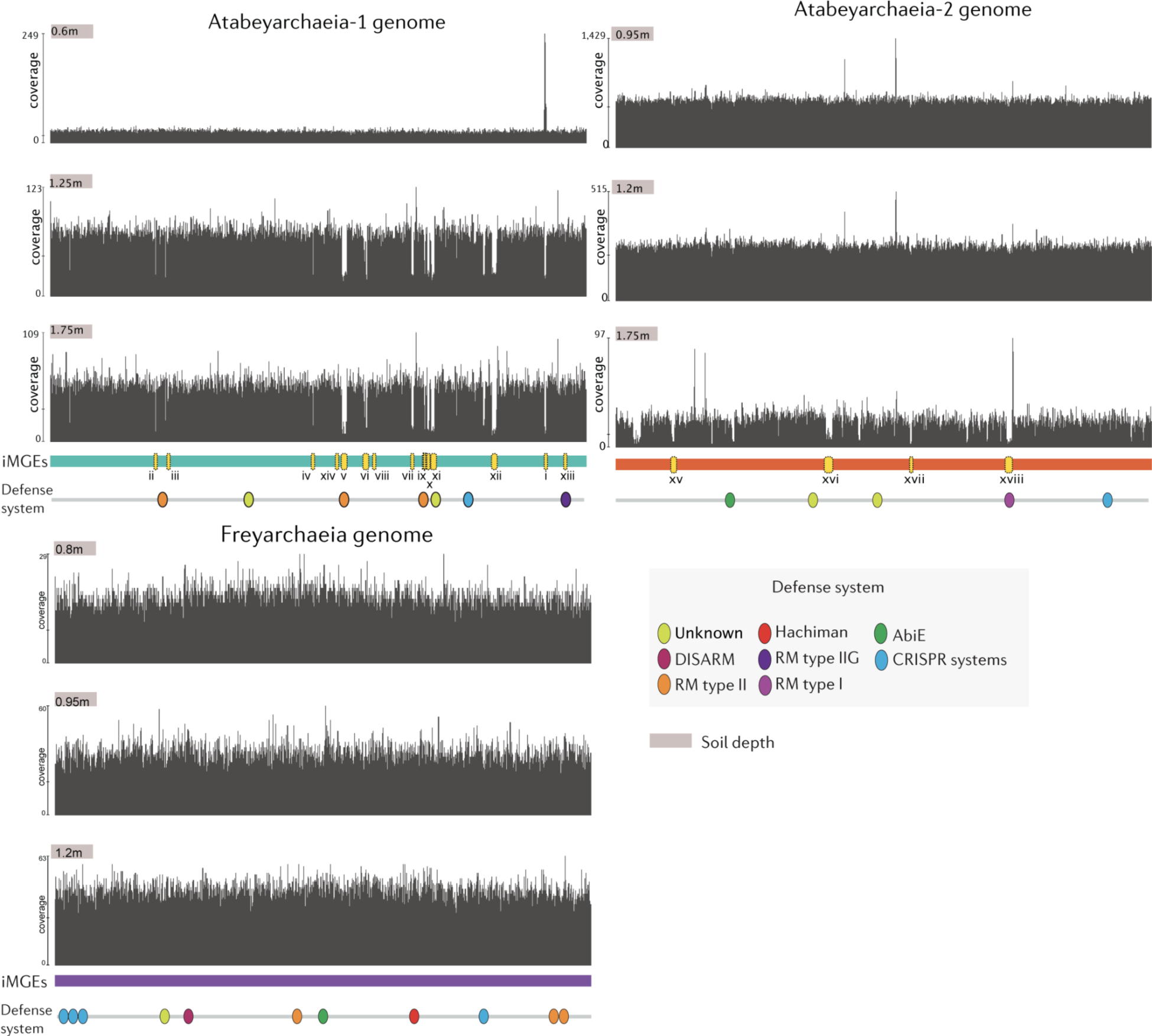
Chromosomally integrated genetic elements and defense systems in soil Asgard Archaea genomes. Each panel depicts the coverage across each complete genome, as determined by mapping to metagenome reads derived from three different soil depth profiles. Regions exhibiting low coverage suggest strain variations associated with specific soil depths and may indicate the presence of integrated genetic elements in only a subset of cells. Notably, the Freyarchaeia genome exhibits even coverage using reads from all sampling depths, with no discernible integrated mobile genetic elements identified. Some of the low-coverage regions are not labeled as potential MGEs, these regions are strain variants with sequences so divergent that read mapping is precluded. Oval symbols indicate predicted defense systems.

Of the 18 integrated MGEs in Atabeyarchaeia, five were classified as insertion sequence-like transposons (IS), three as putative integrative conjugative plasmids of 7.9 - 12 kbp in length carrying integration machinery and cargo genes, two as defense islands, six as pro-viruses, and two could not be classified (**Table S1**). To date, the only Asgard non-viral integrated genetic elements reported are Heimdallarchaeia “aloposons”(Wu et al., 2022), which are transposons that carry cargo genes. These previously reported MGEs do not display any homology at the nucleotide level with those found in Atabeyarchaeia. However, some Atabeyarchaeia integrated genetic elements and these aloposons encode partition proteins (ParB-like) that are distantly related to tyrosine-like integrases (**Figure S1-A**). The Atabeyarchaeia tyrosine-like integrases are most closely related to those found in genomes of Njordarchaeales, Bathyarchaeia and Aenigmarchaeota, which share a similar ecological distribution in terrestrial wetlands and also occur in deep ocean sediments(Seitz et al., 2019) ( **Figure S1-B**).

Ten of the integrated genetic elements coexist in circularized form with their integrated versions in the same metagenomic samples (e.g., **Figure 1**). One of these of particular interest is Atabeya-1 MGE-i (Yucahu-i, in homage to the son of the Taíno goddess Atabey—reflecting our previous designation of the host archaeon as Atabeyarchaeia)—for which the coexisting circularized version in 60 cm deep soil is four times more abundant than the integrated version (**Fig. 2A**). Some of the reads span the genome indicate that a subset of Atabeya-1 genomes lack or have excised this integrated element (**Figs. 1, 2B**), enabling us to determine the exact length of the Yucahu-i to be 10,867 bp. The MGE is inserted following an AATTAACTTAT sequence that is also present at the end of the integrated Yucahu-i and occurs within the excised, circularized version. This region likely represents the attachment (att) site, a unique location within the genome. The low GC content (9%) compared to the genome-wide average (∼50%), suggests that the DNA in this area may exhibit increased susceptibility to cleavage during processes such as excision or integration.

The Yucahu-i element includes 11 open reading frames (**Figure 2A**). Some of the gene products could be functionally annotated using protein homology and *in silico* structural prediction. The first gene encodes a tyrosine recombinase/integrase that likely recognizes and cuts at the AATTAACTTAT motif in the genome and in the circularized version (resulting in Yucahu-i linearization), and may be involved in integration of the linear sequence. The subsequent gene is a Holliday junction resolvase, which likely acts in conjunction with the integrase. We are uncertain if a host integration factor is required, but it is possible that two of the following genes predicted to encode DNA-binding proteins may have this function. Yucahu-i also encodes a superfamily 3 (SF3) helicase that may unwind the DNA and initiate plasmid replication(Guo & Huang, 2010), and a Type II-G restriction-modification (IIG RM) protein fusion that combines endonuclease and methyltransferase activities. Phylogenetic analyses place the IIG RM sequence shares its most recent common ancestor with sequences found in DPANN archaeal genomes (>60% amino acid identity). Basal to this clade are many sequences from bacteria, which supports the inference that the origin of the sequences in question is likely archaeal, potentially acquired via horizontal gene transfer (**Figure S2**). Metatranscriptomic data indicate that the IIG RM gene is transcribed (Supplementary Data). Based on predicted protein functions and presence of the excised, circular (copy number up to 4x) mobile genetic element, Yucahu-i is likely a plasmid.

We investigated how frequently Yucahu-i was integrated in, or coexisted in circular form with, the Atabeya-1 genome by systematically analyzing reads from the 20 soil metagenomes that contained this archaeon (**Figure S3, Table S2**). In 11 samples, Yucahu-i is integrated into essentially all Atabeya-1 cells, however, the data indicate substantial variation in presence/absence of the integrated version and in the copy number of the circularized version (**Figure 2, Figure S3, Table S2**). A few reads from the 70 cm deep soil revealed evidence for the circularization of a 2,644 bp element that is integrated within Yucahu-i. We refer to this as mini-Yucahu-i (**Figure 3-C**). Its presence highlights the genetic plasticity of the plasmid. The mini-Yucahu-i carries a putative ParG, a hypothetical protein, and Holliday junction ATP-dependent DNA helicase RuvB. Interestingly, the identical 11 bp Yucahu-i putative attachment motif is also present adjacent to, and within, a 7,848 bp integrated element in the Atabeya-2 genome (iMGE-xvi). However, the genomes share no detectable similarity, and the percentage of identity of the tyrosine integrases are < 25% and they are phylogenetically unrelated (**Figure S4)**.

**Figure 3.**
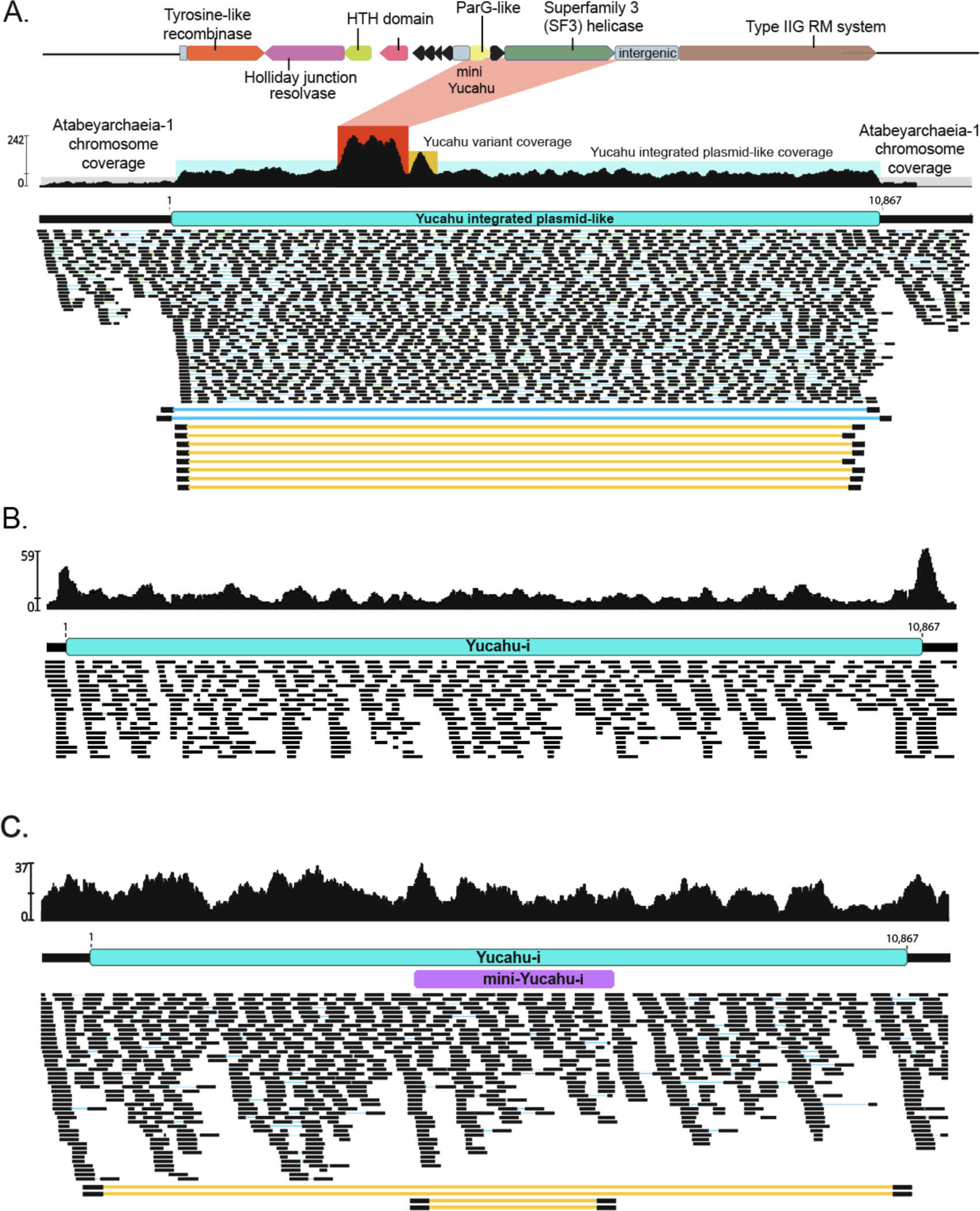
Integration and excision of Yucahu. **(A)** For the 60 cm sample, elevated coverage and paired reads indicate that Yucahu-i is integrated into the genome, excised from some genomes (blue lines) and coexists in circularized form (yellow lines). The red box indicates elevated coverage from a related gene from another genome. **(B)** For the 165 cm sample, low coverage over the MGE and read sequence discrepancies indicate that most cells in this sample lack the MGE. **(C)** For the 70 cm sample, coverage and paired read information indicate that Yucahu-i is integrated into essentially all cells. The circularized Yucahu-i is present but rare. Paired reads pointing out internal to the MGE indicate that a 2,644 bp element has integrated into the plasmid and coexists in circularized form.

### MGEs targeted by CRISPR systems

To explore exogenous MGEs of Atabeyarchaeia and Freyarchaeia, we mined CRISPR spacers from their genomes and matched them to unbinned metagenomic scaffolds from the same wetland soil. More than 30 putative MGE scaffolds are confidently targeted by CRISPR spacers and thus predicted to have once replicated within Atabeyarchaeia. We manually curated them and obtained one complete 20.8 kbp circular plasmid genome, two circular, complete 40 kbp genomes for a pair of closely related viruses, and a circular complete 26.7 kbp genome for an unclassified MGE.

The 20.8 kbp plasmid has 24 open reading frames (ORFs), primarily encoding hypothetical proteins (**Figure S5, Table S3-S4**). It also encodes plasmid proteins such as protein repressor ribbon-helix-helix protein from the copG family (Gomis-Rüth et al., 1998), usually present in bacterial conjugative plasmids. Other predicted proteins are implicated in autonomous replication, such as a DNA primase-helicase and a tyrosine integrase, as well as other genes, involved in nucleic acid processing. Seven of these proteins contain transmembrane domains suggesting the presence of a putative conjugative system or a secretion-like system (**Figure S5**). A protein with a Glu-Glu motif was annotated as an integral membrane CAAX-like protease self-immunity, based on structural modeling and phylogeny (**Figure S6**). It encodes a CTPase with similar function to ParB, a protein typically associated with plasmid chromosome partitioning during replication. Phylogenetic analysis places this protein within a clade that contains MGEs recently discovered in Heimdallarchaeia (**Figure S1-A**). Interestingly, this clade also contains ParB-like proteins from draft genomes of Lokiarchaeia and Thorarchaeia, along with other archaeal genomes (e.g., Sulfulobales), suggesting that this plasmid lineage is widespread in other Asgard archaea. Also included are sequences from *Streptomyces* plasmids. Therefore, these plasmid partitioning genes may have undergone inter-domain horizontal gene transfer.

The circular 40,094 bp virus is predicted to infect Atabeyarchaeia-2 based on CRISPR spacer matches. We named this virus ‘Opia,’ after a mythical creature associated with the Taíno goddess Atabey. Interestingly, we found this virus integrated into the end of a 2.58 Mbp PacBio-derived genome fragment from an related Atabeya-2 strain (Atabeya-2’), confirming its host association. The genome encodes structural proteins such as a capsid-like protein, phage head morphogenesis protein, phage portal protein, tail-like structural proteins and a phage terminase large subunit (K06909:xtmB). Comparison of Opia terminase and capsid proteins with reference archaeal and bacterial virus proteins placed them with those of other Asgard viruses, including Nidhogg, a virus of Helarchaeales (**Figure 4A-B**). The Opia genome harbors genes for various nucleic acid processing proteins, such as a Mu-like prophage protein com and putative transposase, tyrosine recombinase-like, site-specific DNA-methyltransferase, and a ParB-like CTPase. A DNA polymerase sliding clamp subunit (PCNA) like protein (**Figure 4C**) potentially interacts with viral replication proteins, promoting viral DNA synthesis and possibly manipulating host cellular pathways to facilitate viral replication and immune evasion.

**Figure 4.**
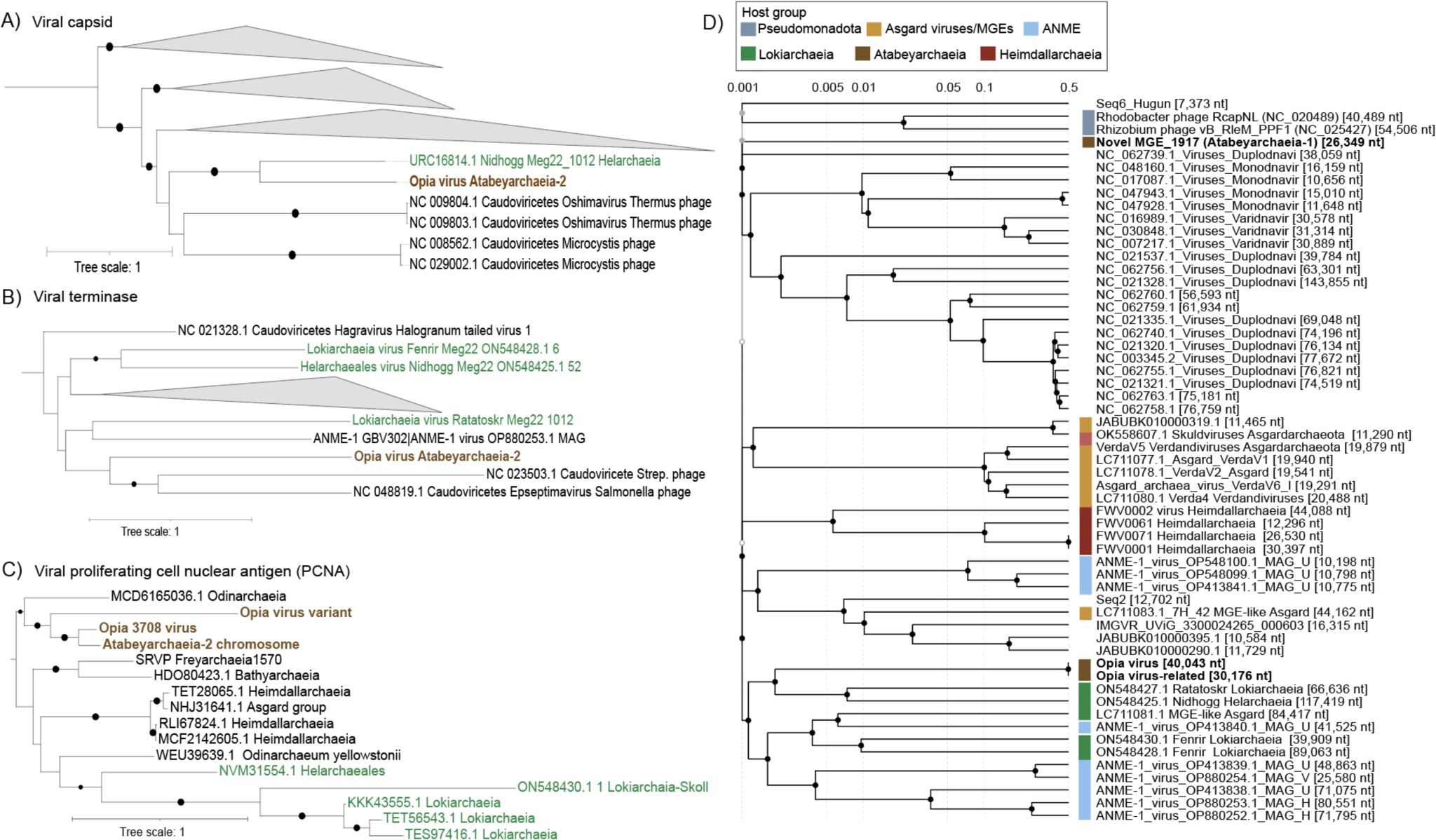
Phylogenetic placement of hallmark structural viral proteins from Opia and whole proteome-based similarity of MGE from Atabeyarchaeia and other Asgardarchaeota. A) Viral capsid B) Viral terminase C) viral proliferating cell nuclear antigen (PCNA) and D) whole proteome-based similarity of MGE from Atabeyarchaeia and other Asgardarchaeota. Boostraps are represented by black circules >80%.

Phylogenetic analysis positioned the Opia PCNA-like sequence within an Asgard archaea clade with a homolog present in the chromosome of Atabeyarchaeia-2. The most similar homolog was a predicted protein from an Njordarchaeales (MCD6165036.1) from the Auka vent field (Speth et al., 2022), suggesting that viruses related to Opia integrate into other Asgard genomes. Homologs of this protein occur in the Sköll viral genome that infects Lokiarchaeia (Medvedeva et al., 2022; Rambo et al., 2022; Tamarit et al., 2022) as well as other archaeal viruses (Mizuno et al., 2019; Raymann et al., 2014). At the whole proteome level, Opia has similarity to the Ratatoskr, Nidhogg, Skoll, and Fenrir viruses (Rambo et al., 2022), known to infect various Asgard archaea (**Figure 4D**).

We identified at least seven distinct Opia virus variant genotypes. The sequences align near-perfectly over >85% of the genomes (**Figure 5A**). All Opia variants (and their identifiable fragments) are exactly targeted by one CRISPR spacer present in loci of both Atabeya-2 and Atabeya-2’ (two identical sequential spacers in Atabeya-2’), despite the presence of the Opia provirus in the Atabeya-2’ genome. The provirus was only partially recovered, so it is impossible to say whether the integrated version differs in the targeted region. A hotspot in the Opia virus genomes encodes a series of genes that are distinctly different in some variants. Included are up to six small cysteine-rich proteins, many with predicted double Zn-binding domains (**Figure 5A**). The genes for specific cysteine-rich proteins occur in different combinations from different genotypes (e.g., one has sequence types A, B, D another A, C, D, and another, C, E). In addition, a three-gene block (one of which has sequence variants) and adjacent intergenic sequences are variably present/absent. Finally, different versions of ParB-like partition proteins occur in the variable region and some lack a C-terminal endonuclease domain (**Figure 5**).

**Figure 5.**
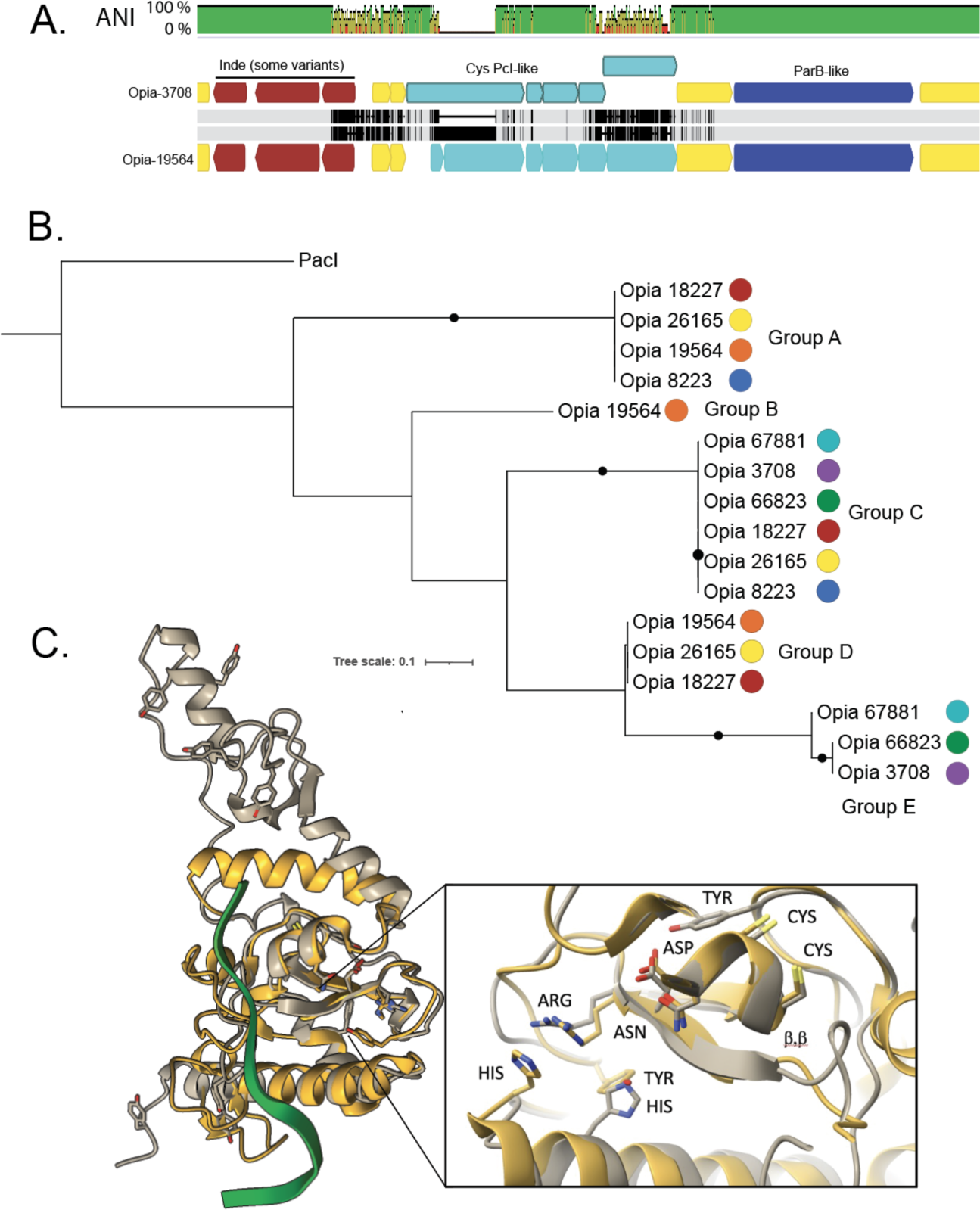
Sequence variation in the cysteine-rich proteins of Opia viruses and comparison to Pacl, a rare-cutting HNH restriction endonuclease. **A.** The aligned ∼5.6 kbp variable region of two Opia viral genotypes. Light gray bars indicate perfect nucleotide identity and thin vertical black lines indicate SNPs. The three genes labeled in brown are present in both Opia-3708 (top) and Opia-19564 (bottom) but are absent in some other geno**type**s. Light blue genes are cysteine-rich (For 3708L, 10, 5, 9, 5, 10 and for Opia-19564, 3, 14, 5, 9, 4, 10 cysteines per protein). Dark blue genes encode ParB-like proteins that are very divergent between some genotypes (Opia-3708 vs. Opia-66823, 67881). Before the 3-gene indel and after the ParB-like gene, the ∼35 kb regions of all genomes are essentially identical. **B.** Phylogenetic tree including the 17 larger cysteine-rich proteins from the variable regions of 7 Opia genomes (numbers represent genome names; for context, see supplementary data. Proteins with identical sequences occur in five different combinations across the genotypes. **C.** Comparison of the active site region of PacI (with 9 cysteines) and the structure of Opia-19564 protein 16 (silver, with 10 cysteines). Active site residues of PDB 3m7k (gold) are displayed based on Figure 1 of Shen et al. (2010). The critical ββα-metal motifs are well aligned. Tyrosine and histidine residues exist in proximity to the active site, although their locations differ somewhat, possibly due to fold inaccuracy.

We predicted the structures of the largest cysteine-rich protein from Opia-3708 (gene 46) and two Opia-19564 proteins (genes 16, 18). All three represent different protein sequence clusters but they share core structural components, including two sets of four cysteines, and an alpha helix in proximity to paired antiparallel beta strands (i.e., ɑ, ββ,-metal; **Figure 5C**). HHpred predicts the Opia-3708 protein (197 aa) to be related to an HNH endonuclease and the best match for the three-dimensional structure (PDB 3M7K) is the Rare-Cutting HNH Restriction Endonuclease PacI, a homing endonuclease that is one of the smallest restriction endonucleases known (142 aa)(Shen et al., 2010). Zinc bound by the four cysteines is required for the DNA cleavage by PacI endonucleases. The H, DR, and CxxCN catalytic residues of 3M7K HNH endonuclease are generally conserved in the Opia proteins (e.g., Opia-19564_16, **Figure 5C**). However, the expected tyrosine residue precedes rather than follows the DR motif and its placement in the predicted structure is offset from that in 3M7K. The histidine active site residue is also slightly differently positioned (**Figure 5C)**. These discrepancies may be attributed to uncertainties in protein folding. However, the positioning of histidine in the location typically occupied by tyrosine suggests its potential involvement in DNA cleavage, as occurs in other HNH endonucleases. Elsewhere, the predicted structures have large regions of positively charged surface, likely involved in DNA binding. These findings suggest that the Opia proteins share characteristics with PacI restriction endonucleases, yet they may represent a novel class of enzymes, likely with homing endonuclease function. (**Figure 5B**). The biochemically characterized PacI homodimer has a target recognition sequence of 5’-TTAATTAA-3’, and cleaves between the internal thymine residues. Pacl endonucleases rely on the absence of the recognition site elsewhere in the host genome. We could not determine the recognition sequence for the Opia Pacl-like homing restriction endonuclease, but apparently it was possible for different combinations of six variants to insert in the same region of a series of Opia genotypes.

We further reconstructed a circular, complete 26,349 bp genome (MGE-9917) for another circularized element that is targeted by three CRISPR spacers from Atabeya-1. MGE-9917 could not be classified as a virus or plasmid based on predicted protein functions. All MGE-9917 genes are encoded on the same strand. The genome features a CT-rich intergenic tandem repeat region with 6, 7, 8 or 9 units of 8 bp in length (variants identified using mapped reads). MGE-9917 encodes at least 14 proteins with transmembrane domains, two of which are 1,402 and 1,202 amino acids in length and lack related sequences in the NCBI database (**Table S4)**. The genome contains a protein that combines an N-terminal ParB-like nuclease domain with a C-terminal S-adenosylmethionine-dependent methyltransferase domain. Additionally, MGE-9917 includes genes for a tyrosine recombinase, a transposase, and a Type IV methyl-directed restriction enzyme featuring an HNH motif. A family of related elements occurs in virtually all of the deep soil samples. One version differs due to the presence of a transposase that is related to those found in the Opia viruses.

### Defense systems and epigenetic regulations in Atabeyarchaeia and Freyarchaeia

We used DefenseFinder (Tesson et al., 2022) and PADLOC(Payne et al., 2022) to identify ten defense systems in Freyarchaeia, five in Atabeyarchaeia-1, and six in Atabeyarchaeia-2. Freyarchaeia harbored at least four different defense system classes, including Type I-B, III-A, and III-D CRISPR-Cas systems, Type II restriction-modification systems, the Hachiman antiphage defense system, and the anti-phage system, DISARM (defense island system associated with restriction-modification), which has not previously been found in Asgard archaea (**Figure S7-A, Table S5-S6**). The DISARM system comprises *drmABC*, a methyltransferase (*drmMI*, N6 adenine-specific methyltransferase or *drmMII*, C5 cytosine-specific DNA methyltransferase), and *drmD* or *drmE*) (Ofir et al., 2018). The Freyarchaeia system includes *drmA*, *drmB, drmC, drmMII*, *drmE,* and drmD (helicase similar to the RNA polymerase (RNAP)-associated SWI2/SNF2 protein) that is present in, classifying this system as a DISARM class II (DISARM-II). Interestingly, *drmD* is a homolog typically found in DISARM class I. The DISARM methylase modifies host CCWGG motifs to distinguish its own DNA from foreign DNA. A specific conformation of the DrmAB complex (trigger loop) inhibits the complex to prevent an autoimmune response (Bravo et al., 2022). DrmA is responsible for DNA targeting in DISARM through multiple non-specific interactions with the DNA backbone (Bravo et al., 2022). By not requiring a specific sequence for DNA binding, DrmA distinguishes this defense system from other common restriction-modification systems, endowing DISARM with a broad spectrum of action against viruses (Tesson et al., 2022). Phylogenetic analysis of the helicase DrmA places the gene within the Euryarchaeia and Chloroflexota, suggesting that this system has been laterally transferred **Figure S7-B.** The Hachiman system is encoded by *hamA*, a hypothetical protein (DUF1837), and *hamB*, a helicase (pfam00271). The molecular mechanism of this system is still unknown (Doron et al., 2018). One potential hint is the presence of an ATP-dependent endonuclease is an OLD (overcoming lysogenization defect) upstream of the *hamAB* locus.

We used DNA polymerase kinetics from Pacific Biosciences (PacBio) metagenomic sequencing data and the Restriction Enzyme database (REBASE) to illuminate the DNA methylation patterns and methylases in the genomes of Freyarchaeia, Atabeyarchaeia-1, and Atabeyarchaeia-2. In the genome of Atabeyarchaeia-1, we identified 13 methylation sequence motifs, of which seven were directly linked to a specific methylase gene. Similarly, Atabeya-2 has 11 methylation motifs, 5 of which could be linked to a methylase. Freyarchaeia has only five detectable methylation motifs (**Table S7A-B**). Interestingly, one of those motifs is CCWGG, which has been characterized as a motif targeted by DISARM class II.

Atabeyarchaeia-1 and Atabeyarchaeia-2 both have 4-methylcytosine (m4C) and 6-methyladenosine (m6A) methylation. Atabeyarchaeia-1 Yucahu-i MGE encodes a Type IIG restriction-modification system that targets a m6A methylation motif and is the only candidate enzyme that could methylate the Atabeyarchaeia-1 genome. The Yucahu-i system is analogous to the MmeI family, which typically recognizes a 6-7 bp motif with adenine as the penultimate base (Morgan et al., 2009). The motif GYATGAG (m6A) was methylated at 66% of sites within the Atabeyarchaeia-1 genome and could represent the active methylation motif of this RM system. In contrast, Freyarchaeia has 5 m4C motifs but no m6A methylation motifs.

### Integrated mobile-like regions encode eukaryotic signature proteins

We identified two small GTPases in a region of the Atabeyarchaeia-1 genome that appears to be enriched in genes often associated with mobile genetic elements. This region is absent in some Atabeya-1 strains. The classification of these proteins as GTPases is supported by sequence and structural homology, as well as, structural predictions in comparison to reference eukaryotic sequences (**Supplementary Text**). Based on phylogenetic analysis, these proteins cluster with the unannotated Arf GTPases, ArfX, found in single-cell eukaryotes *Naegleria gruberi*, *Spironucleus salmonicida, Gefionella okellyi*, and Arfrp 1b in *Pygsuia biforma* (**Figure S8**).

## Discussion

We identified chromosomally integrated and coexisting MGEs that replicate in Atabeyarchaeia by leveraging extensive sequencing of a series of soil samples where soil depth and biogeochemical conditions select for different strain variant populations. By examining read mappings across multiple samples from the same environment, we established a larger repertoire of MGEs than could be found from the analysis of any single metagenome. Integrated elements and coexisting circularized MGEs range from 2.5 to 40 Kb in length; all complete MGEs were circular and at least some replicate bidirectionally. We could not confidently identify MGEs by changes in read mapping abundances in the Freyarchaeia genome. This might indicate either stable integration into the Freyarchaeia genome across the entire population studied (thus excision sites and coexisting versions of circular elements were not detected by our methods) or that, the degree of association with MGEs may vary dramatically between Asgard archaeal lineages.

The presence of coexisting integrated and free, circularized MGEs, likely mostly plasmids, as well as variation in copy number of circularized elements and in the fraction of cells with integrated elements suggest regular movement of MGEs into and out of the Atabeyarchaeia chromosomes. Insertion/excision and variation in copy number may enable Atabeyarchaeia to respond to changes in their environment. For example, MGEs may behave symbiotically, and increase in MGE copy number (thus gene content) serves as a response to increased pressure from other MGE(Krupovic et al., 2019). The tiny mini-Yucahu indicates another layer of genomic variability, as just this portion of the host MGE can excise.

Interestingly, the attachment motifs for Yucahu-i plasmid-like Atabeya-1 MGE and for a circular, unclassified and essentially unrelated MGE linked to Atabeya-2 are exactly the same, implying that the very different integrases of each (25% aa ID) recognize and cut at the same motif. Protein sequence divergence may enable host chromosome specificity, yet the active site apparently evolved to target the same motif.

The novel cluster of genomically similar Opia viruses of Atabeya-2 and 2’ encode sequential cysteine-rich proteins inferred to have nuclease activity due to their distant homology to PacI. These occur combinatorial patterns (**Figure 5B**) that may have arisen via recent recombination events in which these putative endonucleases may have played a role. The multiple variants might form heterodimers rather than the normal homodimers expected for Pacl, possibly extending target recognition. The results suggest the importance of diverse nuclease activity for these viruses.

The Opia virus proteomes exhibit similarities (e.g., capsid, tube and tail proteins)to those of tailed viruses, which commonly replicate in bacterial and archaeal hosts from hypersaline environments (Senčilo & Roine, 2014). They are quite distinct from those of eukaryotic viruses, supporting the suggestion that, despite the evolutionary relationship between Asgard archaea and eukaryotes, their viruses display no obvious evolutionary relationships (Medvedeva et al., 2022; Rambo et al., 2022; Tamarit et al., 2022).

Genome context (dominated by MGE-associated genes) and apparent excision of the region encoding the GTPases from some strain genotypes supports the inference that some Atabeyarchaeia MGEs encode eukaryotic signature proteins. ARF GTPases are involved in membrane trafficking in eukaryotes and have been previously described as ESPs in Asgard archaea (Eme et al., 2023; Spang et al., 2015). The presence of these GTPases on putative MGEs provides the first indication that increase in cellular complexity could be associated with the transfer of eukaryotic signature proteins via MGE (**Figure S8, Figure S9, Table S8)**.

Our results suggest several examples of genes integrated into Atabeyarchaeia genomes or in their coexisting MGEs that have nearest homologs in the bacterial domain (e.g., ParB, DrmA, and Type IIG restriction-modification enzymes). These findings are consistent with recent work on cross-domain gene transfer via movement of integrons associated with diverse MGEs (Ghaly et al., 2022) and extend earlier work inferring the acquisition of archaeal genes by bacteria (e.g., Hug et al., 2013). As we discover new Asgard archaea from genome-resolved metagenomes, we can expect to find further parallels between bacterial and archaeal immune systems. These associations could have implications for the evolution of eukaryotic immune systems (Wein & Sorek, 2022).

Type IIG restriction-modification systems, to our knowledge, have not previously been associated with archaeal MGEs. The observation that the only methylase seemingly able to methylate at the host genome’s well-represented m6A motif is carried by Yucahu-i, and that it is transcriptionally active, suggests that the Yucahu-i Type IIG plays a significant role in host genome epigenetic modification. This MGE-encoded system may protect its host Atabeyarchaeia against infection by other MGEs. This behavior aligns with the emerging perspective that defense systems themselves can serve as mobile genetic elements (Rocha & Bikard, 2022; Wu et al., 2022).

To our knowledge, these are the first metagenome-derived asgard archaeal complete genomes for which methylation patterns have been reported. PacBio sequences corresponding to these complete, manually curated genomes(Valentin-Alvarado et al., 2023) were used to infer the methylation motifs and to determine the fraction of sites that were methylated, as well as the methylation patterns of their newly reported MGEs. Using REBASE, which features all biochemically characterized methylases, it was possible to link methylated sites with likely methylases encoded on both genomes and MGEs. Relatively little is known about genome methylation in archaea, especially in Asgard archaea(Anton & Roberts, 2021). These genomes and their methylation motif data presented here provide a starting point for detailed biochemical studies to expand the known inventory of archaeal methylases. The higher number of methylation motifs in Atabeyarchaeia compared to the Freyarchaeia genome could be an evolutionary response to the larger inventory of mobile genetic elements associated with Atabeyarcheia.

Our results brought to light new MGEs, defense systems, and epigenetic patterns in these soil-associated Asgard archaea. The description and annotation of the first complete Asgard plasmids opens a route toward the development of tools for genetic manipulation of these organisms.

## Conclusions

We leveraged the read diversity inherent to population genomic data, long-read sequencing, methylation pattern analysis, comparative genomics and functional and structural prediction to explore integrated and coexisting genetic elements of one group of Asgard archaea. These analyses brought to light an extensive landscape of mobile genetic elements (MGEs) that associate with Atabeyarchaeia, including viruses, plasmids and as yet unclassified entities. The excision, insertion and changes in copy number of these MGEs may enable adaptation to changing conditions, have contributed evolution, and possibly to the acquisition and spread of genes linked to the origin of cellular complexity. The availability of MGEs that could be adapted for delivery of genome editing tools in a community context (Rubin et al., 2022) may pave the way for genetic manipulation of these archaea.

## Methods

### Sample acquisition, nucleic acid extraction and sequencing

DNA and RNA extractions and shotgun metagenome library construction were described in ref(Al-Shayeb et al., 2022; Valentin-Alvarado et al., 2023). Briefly, we collected soil cores from a seasonally flooded wetland (SRVP) in Lake County, California, in October 2018, October 2019, November 2020 and October 2021 (38°41’39”N 122°31’36”W 571m). Samples were frozen in the field using dry ice, and kept at -80 C until extraction. The Qiagen PowerSoil Max DNA extraction kit was used to extract DNA from 5-10 g of soil, and the Qiagen AllPrep DNA/RNA extraction kit was used to extract RNA from 2 g of soil. Samples were sequenced by the QB3 sequencing facility at the University of California, Berkeley on a NovaSeq 6000. Read lengths for the 2018 DNA samples and the RNA samples were 2x150 bp and 2x250 bp for the 2019-2021 DNA samples. A sequencing depth of 10 Gb was targeted for each of 2018, 2020, and 2021 samples, and 20 Gbp for each of the 2019 samples. PacBio sequencing was obtained from a subset of deep soil samples from 2021 via the University of Maryland sequencing facility. Samples from September 9, 2021 from 140cm and 75cm were sequenced using a Sequel II to generate PacBio HiFi reads. Reads were quality trimmed using BBDuk (bbduk.sh minavgquality=20 qtrim=rl trimq=20)(Bushnell, 2014)and assembled with hifiasm-meta(Feng et al., 2022).

### Discovery of integrated genetic elements using closed complete genomes

Our manual approach to identifying integrated MGEs was based on 1) anomalously low or high coverage over a region, 2) short reads supporting the excision (see **Figure 1**), and 3) genomic architecture and gene annotations. To investigate the integration and excision state of the Yucahu plasmid, we employed a combination of sequencing and bioinformatic analyses. The metagenome reads were aligned to the reference genome sequences. Read coverage was employed to determine the relative abundance and distribution of each MGE across the different samples. Additionally, in-depth examination of the reads allowed for the identification of integration events, mini elements, and Mobile Genetic Elements (MGEs) within the plasmid. To further characterize the MGE, we conducted comparative genomics analysis, comparing genomes lacking the MGE with those containing it. This enabled us to accurately determine the length of the MGE and pinpoint the specific sites of integration within the plasmid.

### Identification and genome curation of Atabeyarchaeia-associated exogenous MGEs

We used metagenomic datasets to search for candidate mobile genetic elements associated with Atabeyarchaeia and Freyarchaeia. Screening was based on taxonomic profile, GC contents and CRISPR-based targeting (see below). All the candidate contigs were manually curated using Geneious Prime-2023.1.2. The manually curated genomes were de novo reconstructed from high-quality Illumina metagenomic data. Manual genome curation methods generally follow(Chen et al., 2020). Long-read PacBio data were used to verify and expand the sequence dataset. Replichores of complete genomes were predicted according to the GC skew and cumulative GC skew calculated by the iRep package (gc_skew.py) Complete MGE genomes with viral structural genes were classified as viruses and genomes that did not have viral structural genes were categorized as plasmid or other genetic elements such as transposons, conjugative elements, and unclassified.

### CRISPR-Cas systems and classification of soil Asgard-associated viruses

CRISPR-Cas systems in Atabeyarchaeia and Freyarchaeia genomes were identified using CRISPRCasTyper v1.8.0(Russel et al., 2020). Spacers were extracted from reads by mapping reads to the corresponding CRISPR arrays via BBMap (sourceforge.net/projects/bbmap/). Recruited spacers were matched against all assembled scaffolds with ≤1 mismatch using Bowtie v1.3.1(Langmead et al., 2009). Scaffolds that are targeted by CRISPR spacers and not affiliated with microbial genomes were curated manually to completion. The phylogenetic classification was predicted based on genome-wide similarities using ViPTree whole proteome-based similarity of MGE from Atabeyarchaeia and other Asgardarchaeota (**Table S9**).

### Coverage calculation of integrated genetic elements and host-chromosome

We aligned metagenome reads to reference genomes using the BBMap’s short-read aligner. A minimum identity threshold of 0.95 was required, and ambiguously mapping reads were discarded. The coverage values of the MGEs, and total genome coverage values were calculated from the alignment/map (BAM) files. The positions of integrated elements were defined. For the genome coverage calculation, the entire genome was divided into two sections: the region before the start of the larger integrated element and the region after the end of the larger integrated element. This approach enabled systematic, efficient calculation of the coverage of integrated elements within genomes.

### Methylation analysis via REBASE and Single Molecule, Real-Time (SMRT)

Methylation patterns within the genomes of Atabeyarchaeia and Freyarchaeia were investigated by mapping PacBio circular-consensus reads metagenomic reads to each of the three curated circular reference genomes for Atabeyarchaeia-1, Atabeyarchaeia-2, and Freyarchaeia using minimap2. The resulting BAM files were then processed and analyzed to identify methylation patterns using the ipdSummary and motifMaker commands in the SMRT Link analysis software package (v11.0; Pacific Biosciences, Menlo Park, CA, USA). To annotate methyltransferase (MTase) activities and restriction enzyme sites, the sequenced genomes and identified methylated motifs were compared against the Restriction Enzyme Database (REBASE)(Blow et al., 2016; Roberts et al., 2015). This comparison enabled the annotation of methylation sites and the determination of specific motifs associated with methyltransferase activity and restriction-modification systems within these archaeal genomes.

### Phylogenetic analysis of hallmark proteins present in MGEs

We compiled the top 25 to 50 best matches for candidate proteins from the NCBI database, ggKbase and UniProt. The sequences were aligned using MAFFT with the parameters –localpair --maxiterate 1000 --reorder. Following alignment, each was trimmed using TrimAl, applying a gap threshold of 0.7. The final alignments underwent manual inspection using Geneious (see above). Maximum likelihood trees were inferred using IQ-TREE version 1.6.12, employing the auto option for model selection, a bootstrap value of 1000, and identifying the best-fit model for constructing the final trees. The details of all models used are included in the description of each figure. Trees were visualized using iTOL. All hallmark MGEs protein alignments and trees have been provided in the supplementary data for further reference.

### Opia virus proteins structure predictions

Proteins were structurally modeled using AlphaFold2(Jumper et al., 2021). Foldseek (van Kempen et al., 2023) was employed to identify structural homologs and structural alignments and comparisons to the modeled proteins were conducted using UCSF ChimeraX (Meng et al., 2023). Active site contents were derived from published descriptions for the reference structures.

### Structural analyses and structural phylogeny of the ESP

The protein sequences of small GTPases found were analyzed along with eukaryotic small GTPases identified as sequence homologues (**Figure S7-8**). These sequences were submitted for structural modeling using ColabFold v1.5.240 (Mirdita et al., 2022). Multi-sequence alignments were performed using the MMseqs2 mode and the AlphaFold2_ptm models. Two recycling steps were employed to improve model prediction. The structural models were used as queries to search for structural homologues in the RCSB Protein Data Bank using FoldSeek easy-search feature, with a cutoff of >15% identity and <E-05 e-value.

Protein structures identified by FoldSeek were integrated with the models generated in ColabFold. A multi-structural alignment (MSTA) of these structures was carried out using the default parameters of mTM-align (Dong et al., 2018) (**Figure S7-C**). The resulting MSTA was further analyzed using IQ-TREE v2.0.3 (Model LG+I+G4 chosen according to BIC), yielding the dendrogram in **Figure S7-B**. The pairwise matrix obtained from the mTM-align process (**Table S8**) was utilized to select proteins suitable for 3D reconstruction of their alignments (**Figure S7-D**). This was done using the Needleman-Wunsch algorithm and the BLOSUM-62 matrix within the ChimeraX software (Meng et al., 2023)

## Reporting Summary

## Data availability

Prior to publication, the genomes reported in this study can be accessed via https://ggkbase.berkeley.edu/SRVP_asgard/organisms.

## Author contributions

The study was designed by LEVA and JFB. Samples collection and nucleic acid extractions were performed by LEVA, JFB, ACC, and L-D.S. Metagenomic data were generated by LEVA, RS, ACC and JFB. Genome binning was done by LEVA and JFB. Genome curation was conducted by LEVA, L-D.S and JFB. MGEs genome analyses were performed by LVA and JFB. Correlation analyses were performed by ACC and LEVA. Phylogenetic analyses of hallmark MGE proteins were conducted by LEVA. LEVA AND JFB performed analysis of integration and excision events of MGEs. LEVA and L-D.S analyzed whole viral proteomes. KEA, PL, VDA and BJB performed the eukaryotic signature protein analysis. LEVA, ACC and RJR performed the methylation analyses. RS contributed to bioinformatics analyses. BJB and DFS provided feedback on the study design, methodology and financial support. LEVA and JFB wrote the manuscript, with contributions from all authors

## Acknowledgments

This research was funded by the Bill & Melinda Gates Foundation (Grant Number: INV-037174 to JFB), a University of California Dissertation-Year Fellowship (to LEVA), a Stengl-Wyer Graduate Fellowship and University of Texas Graduate Continuing Fellowship (to KEA), the Innovative Genomics Institute at UC Berkeley. This work was also supported by the Moore-Simons Project on the Origin of the Eukaryotic Cell, Simons Foundation 73592LPI (https://doi.org/10.46714/735925LPI, to (BJB) and Simons Foundation early career award 687165 (to BJB). The findings and conclusions contained within are those of the authors and do not necessarily reflect positions or policies of the Bill & Melinda Gates Foundation.D.F.S. is an Investigator of the Howard Hughes Medical Institute. We thank Dr. Basem Al-Shayeb and Dr. Marie C. Schoelmerich for his contribution to field work and generation of sequence datasets, Dr. Jacob West-Roberts, Shufei Lei and Jordan Hoff for bioinformatics support, and Dr. Mart krupovic and Dr. Gavin Knott for helpful discussion. Dr. Lin LinXing Chen for critial reading over the manuscript.

## Inventory of Supporting Information Document

### Supplementary Text

The ARF GTPase family is a multi-function group that may be involved in processes such as vesicle transport, lipid modification, organelle morphology (e.g., mitochondrial fusion), and adhesion. Phylogenetic analysis indicates that Atabeyarchaeia, Freyarchaeia, and iMGE-xii proteins belong to a group characterized mainly by unannotated Arf eukaryotic representatives. Thus to get insights into their potential function we determined i) their subcellular localization via (PSORTb v3.0 Archaea), ii) structural predictions (AlphaFold), and iii) structure-based phylogenies (**Supplemental Figure 8, Figure S9, Table S3**). Similar to known eukaryotic proteins PSORT identified localization as cytoplasmic or unknown. From the ArfX cluster on the tree, we chose 5 proteins from Atabeyarchaeia (Atabeya-1, Atabeya-2, and Atabeya-3) and Freyarchaeia genomes, the 2 proteins from the MGEs, 4 proteins from Eukaryotes, and 4 additional Eukaryotic proteins from Arl2 and Arl8 as the outgroup for localization and structural analysis. Then, we searched the predicted GTPase structures of the iMGE-xii_6 and iMGE-xii_29 against the RCSB Protein Data Bank and aligned the predicted and validated structures to curate the structural phylogeny (**Figure S9**, **Figure S9B**). Predicted structures of the iMGE-xii proteins showed strong similarity to the predicted GTPase from Freya (RMSD: 1.606), predicted GTPase from *Spironucleus salmonicida* (single-cell flagellate) (RMSD: 2.476), and protein structures obtained by electron microscopy in a *Homo sapien* (RMSD: 2.252) (**Figure S9B**). Filtered structural alignments (>15% identity) of predicted iMGE GTPases to the RCSB Protein Data Bank (PDB) showed matches were >95% and >91% eukaryotic for iMGE-xii_6 and iMGE-xii_29, respectively (**Table S8**)(Berman et al., 2000). The G domain region of the GTPase is composed of 5 α-helices and 6 β-sheets, and the main function is to bind and hydrolyze guanine nucleotides. The small proteins work as biotimers by differentiating between their active and inactive state. Regions called switches induce the conformational changes between the GTP-bound state (active form) and the GDP-bound state (inactive form)(Vetter & Wittinghofer, 2001). The structural alignments showed a conserved G domain and switches of these GTPases (**Figure S10**), suggesting these GTPases act as biotimers, initiating or terminating the associated protein complexes’ activity(Takai et al., 2001).

**Table S1:**
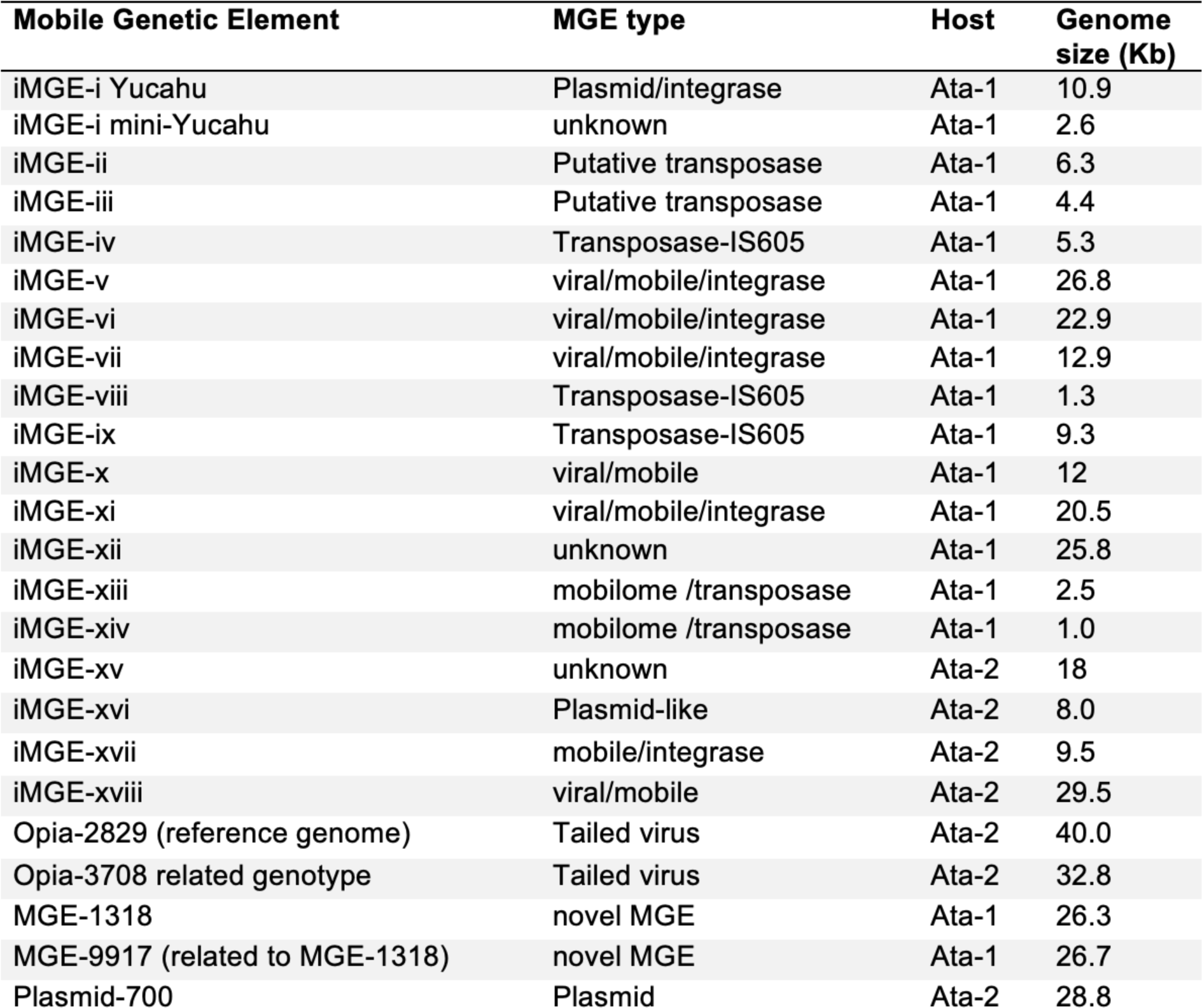
Manually curated genomes for chromosomally integrated genetic elements and extrachromosomal elements. MGE type is the closest classification based on gene content and genome structure. The host was determined based on matches of the MGE squences to spacers from the CRISPR systems.

**Table S2.**
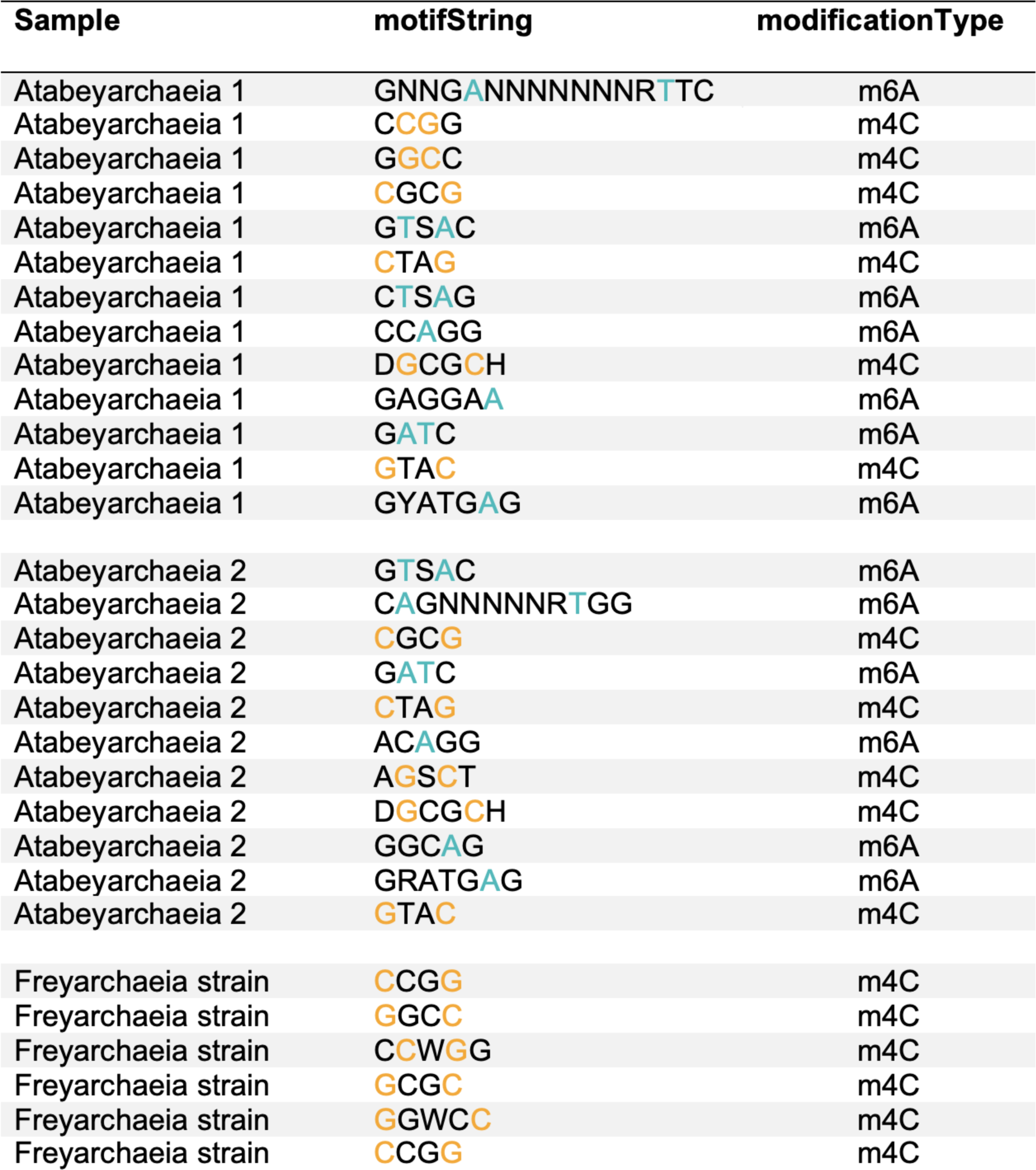
Methylation patterns in Atabeyarchaeia and Freyarchaeia genomes based on PacBio data.

**Figure S1:**
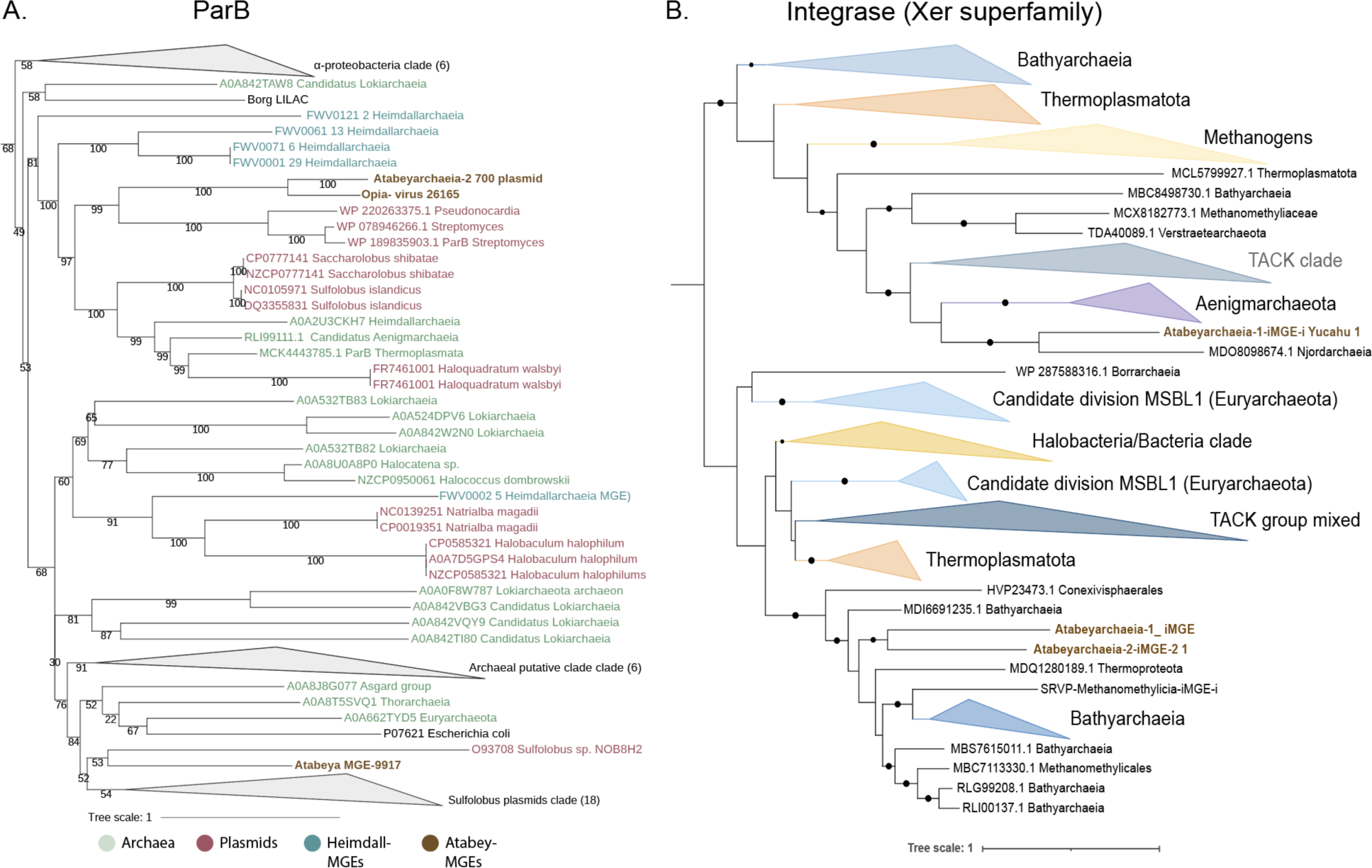
Phylogenetic analyses of ParB and tyrosine-recombinase-like proteins across archaeal mobile genetic elements. **A.** Phylogenetic tree for ParB and ParB-like Proteins. Analysis incorporates reference sequences of *E. coli* ParB (Pfam08775) and *Sulfolobus* conjugative plasmid ParB (NOB8H2), alongside representative ParB-like domains from select archaeal and bacterial genomes. Selection criteria for inclusion involved a rigorous screening of the top 25 protein hits from the NCBI database. Sequence alignments were conducted using MAFFT (v7.310) in ‘auto’ mode, with subsequent optimization by trimAl (v1.4.rev15) applying a 0.7 gap threshold. Phylogenetic trees were initially inferred using IQ-TREE (v1.6.1) under the LG+FO+R model. This segment details **B.** Phylogeny of tyrosine-recombinase-like proteins within integrated genetic elements of Atabeyarchaeia-1 and Atabeyarchaeia-2. The analysis selected the top 25 protein hits from the NCBI database, aligned with MAFFT (version 7.310) on ‘auto’ setting, and refined the alignment with trimAl (version 1.4.rev15) using a 0.7 gap threshold. The phylogenetic framework was constructed using IQ-TREE (version 1.6.1), adopting the LG+FO+R model.

**Figure S2:**
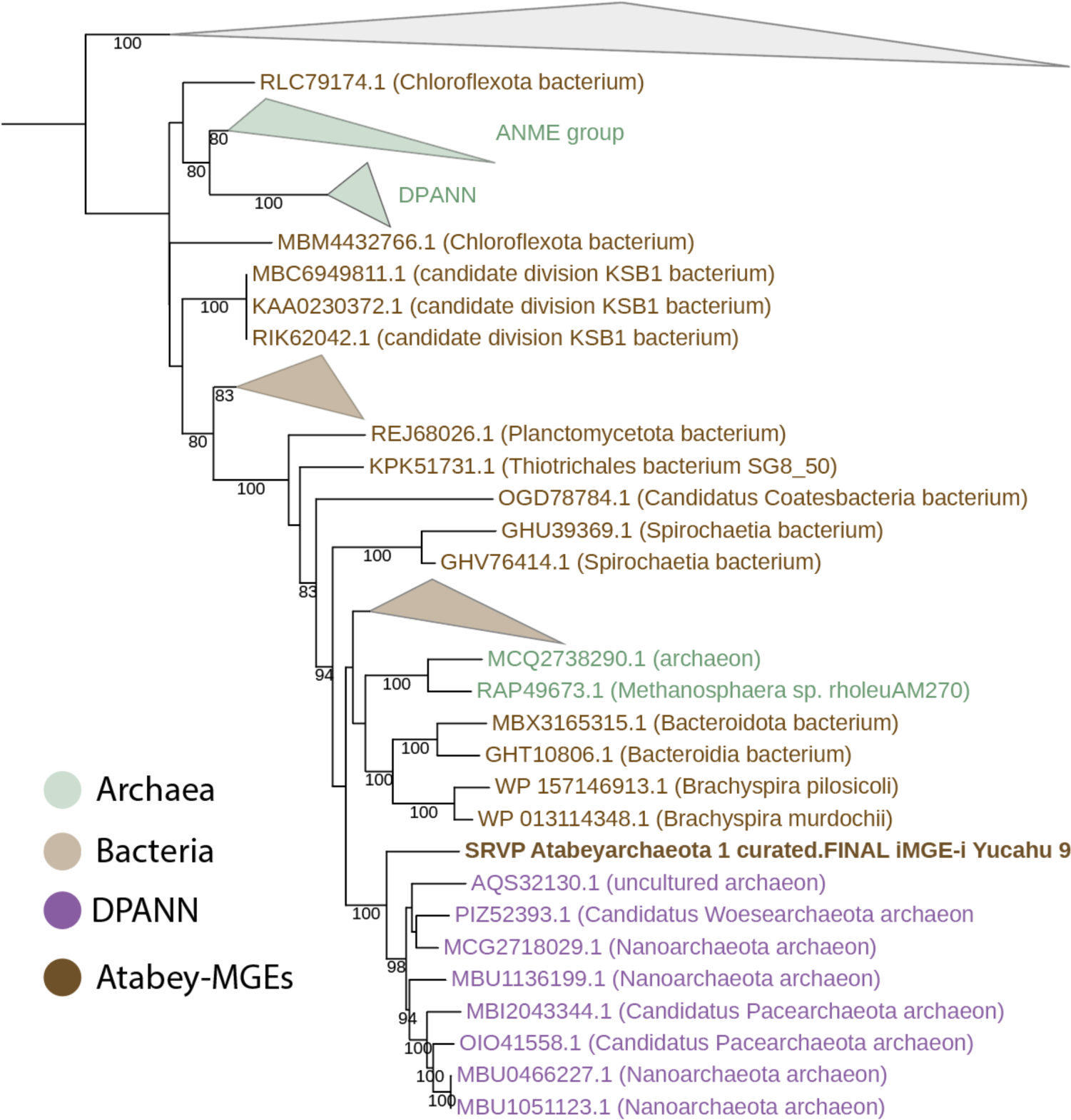
Maximum likelihood phylogeny of Type II-G restriction-modification (IIG RM) protein fusion that combines endonuclease and methyltransferase present in Yucahy MGE. The phylogenetic tree was initially constructed with IQ-TREE (version 1.6.1) employing the LG+FO+R model for the first iterations. Subsequent refinement involved several rounds of manual branch checking to ensure the accuracy and reliability of the phylogenetic relationships depicted.

**Figure S3:**
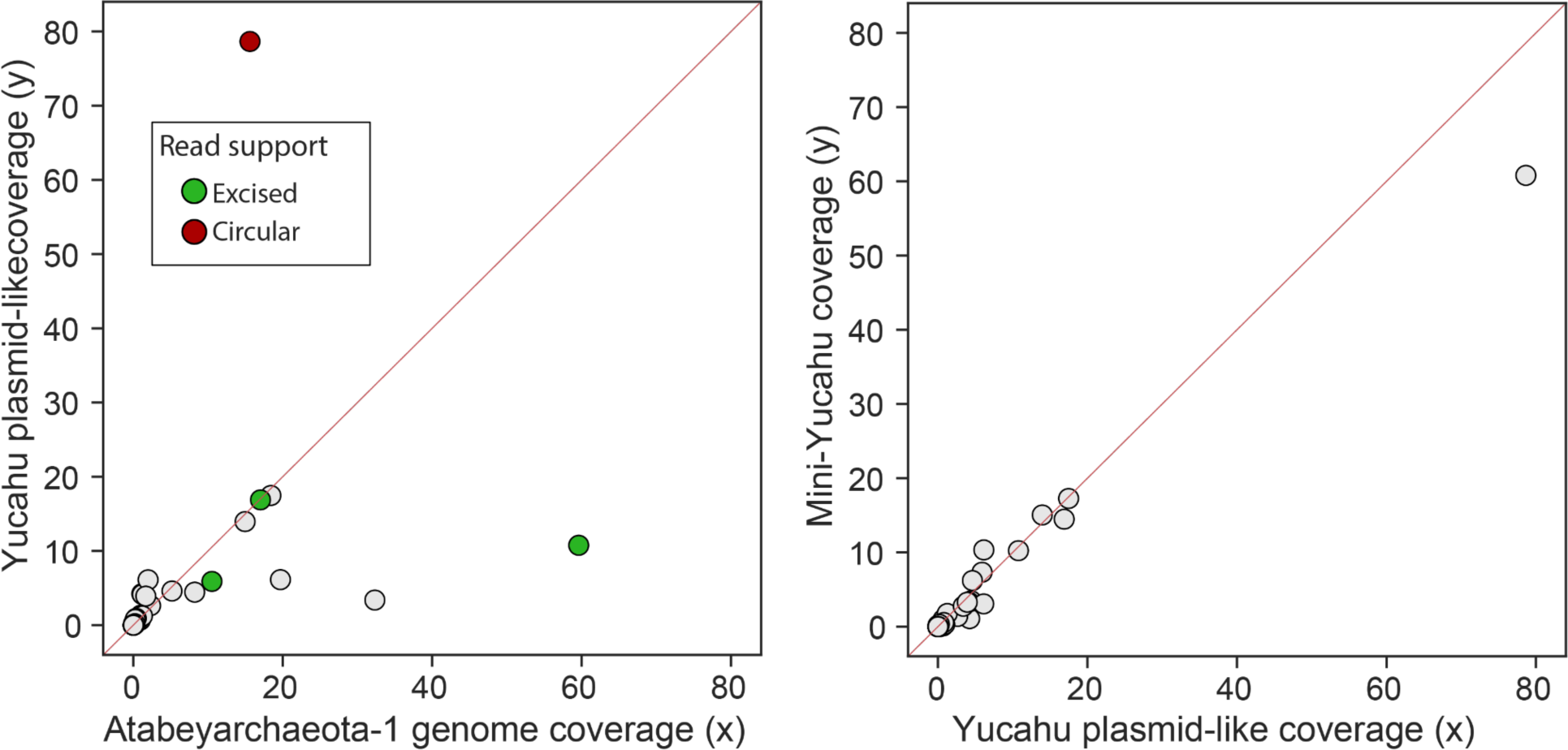
Cross-sample Atabeyarchaeia-1 plasmid integration. **A.** Correlations of the genome coverages between integrated plasmid Yucahu and Atabeyarchaeia **B.** Correlations of the Yucahu plasmid genome and the mini-Yucahu element

**Figure S4.**
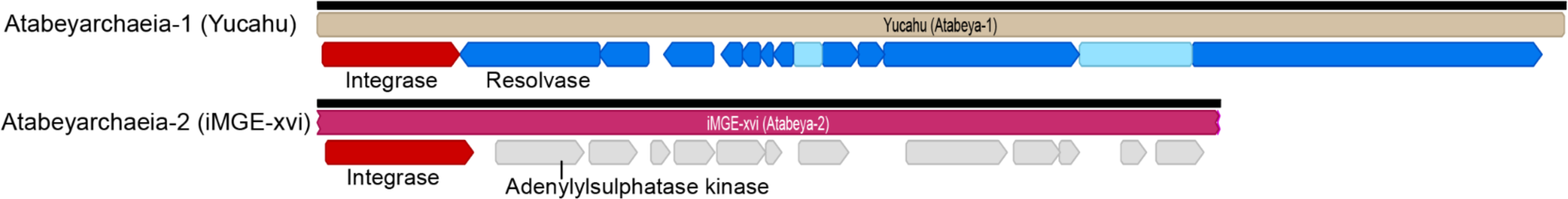
Comparison of gene content of Yucahu-i (Atabeyarchaeia-1), top. and iMGE-xvi (Atabeyarchaeia-2), bottom. Annotated proteins of Yucahu-i include a tyrosine recombinase and a protein with similarity to adenylylsulfate kinases. The iMGE-xvi has 12 open reading frames, most encoding hypothetical proteins or proteins of unknown function. Three proteins are predicted to have 5 - 8 transmembrane domains. One is a putative membrane-bound serine protease of the ClpP class involved in the proteolysis of misfolded and defective proteins(Moreno-Cinos et al., 2019).

**Figure S5.**
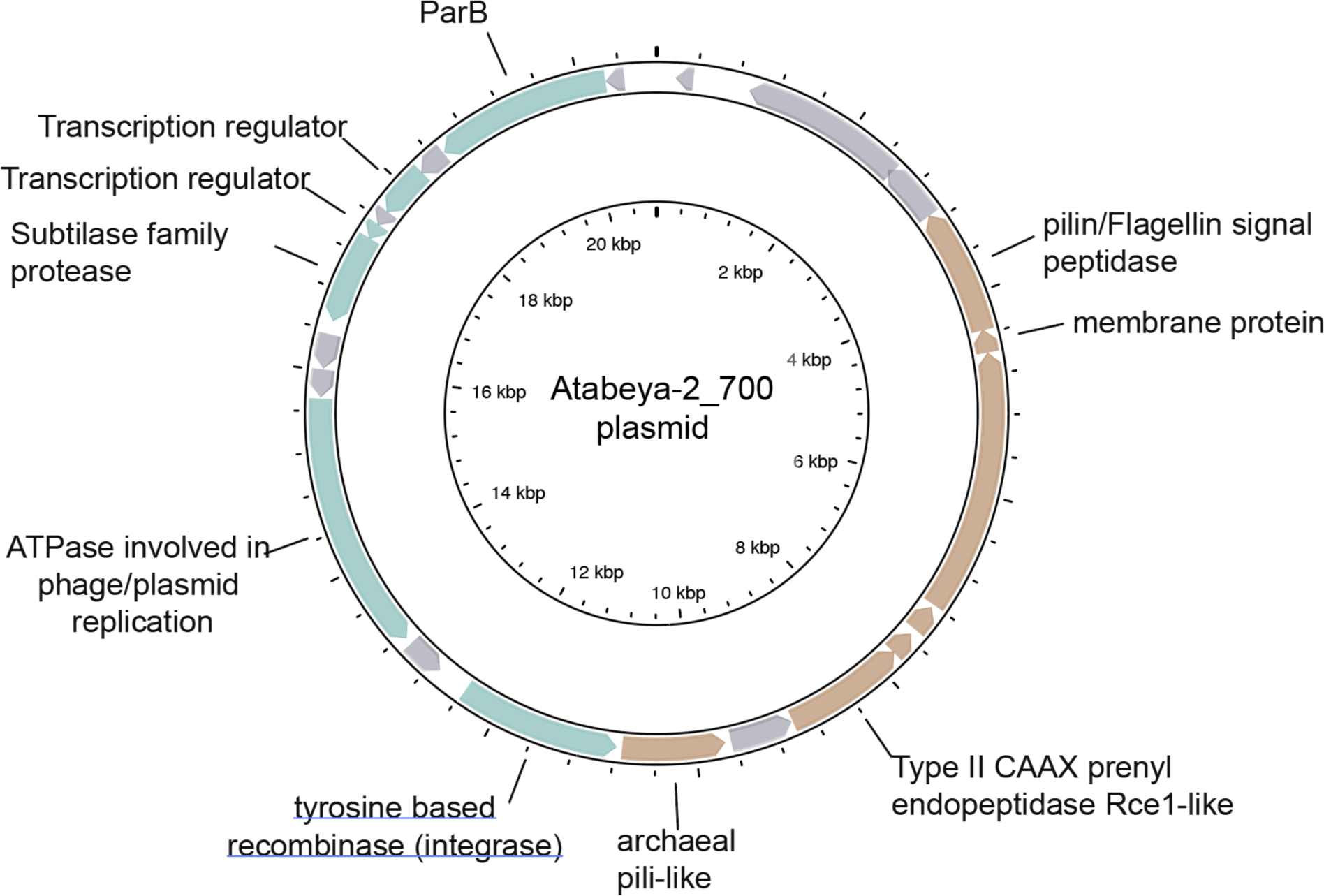
Genome diagram of Atabeyarchaeia-2 putative plasmid scaffold_700. The plasmid has 23 open reading frames and gray-colored genes are hypothetical. Genes colored brown have transmembrane domains and are predicted to be extracellular.

**Figure S6:**
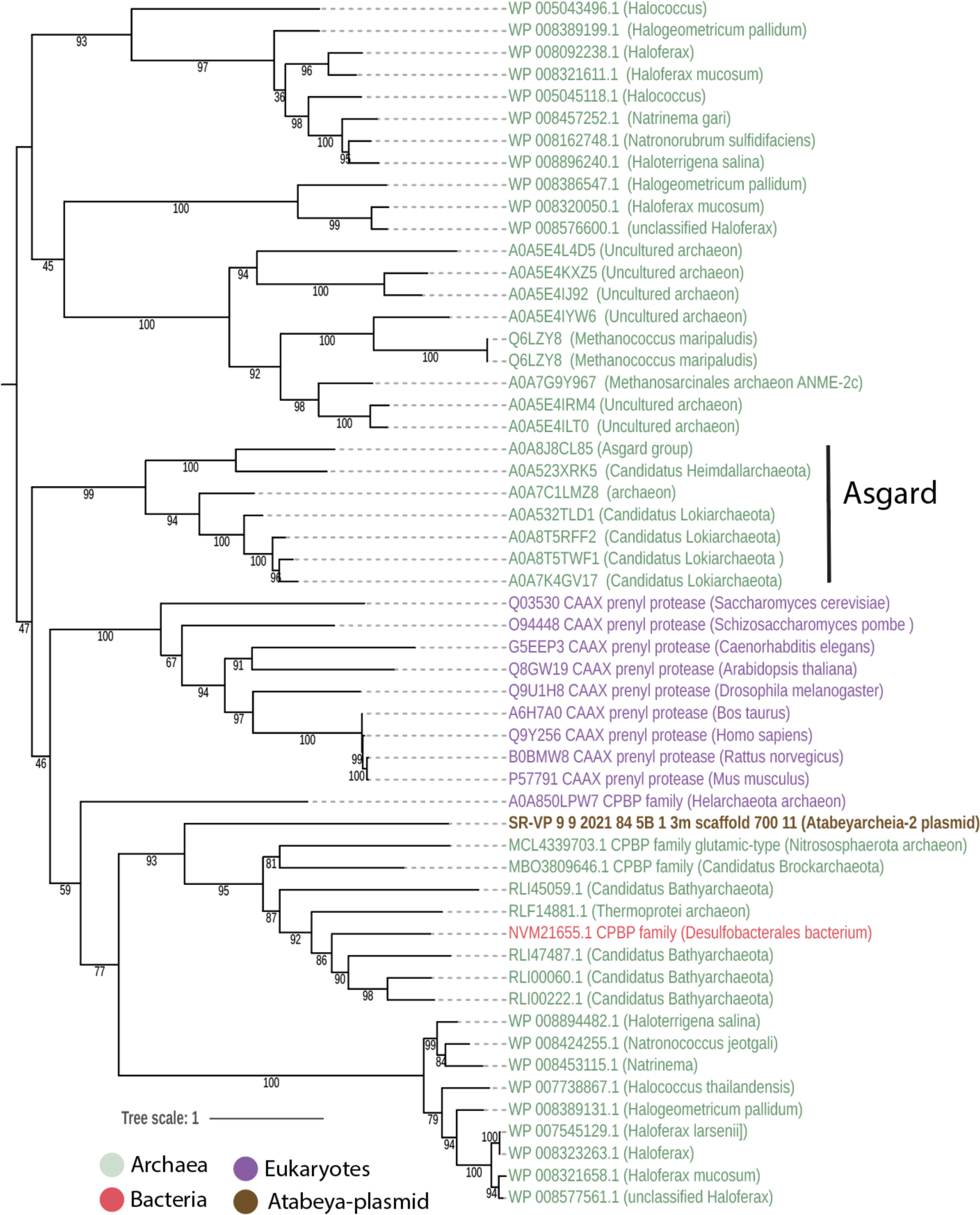
Maximum likelihood phylogeny of CAAX proteases identified in the Atabeyarchaeia-2 plasmid_700. For this analysis, the top 50 protein hits from the NCBI database were selected. Sequence alignment was performed using MAFFT (version 7.310) with the ‘auto’ setting, and the alignment was subsequently trimmed with trimAl (version 1.4.rev15) using a gap threshold of 0.7 (-gt 0.7). The phylogenetic tree was initially constructed with IQ-TREE (version 1.6.1) employing the LG+FO+R model.

**Figure S7:**
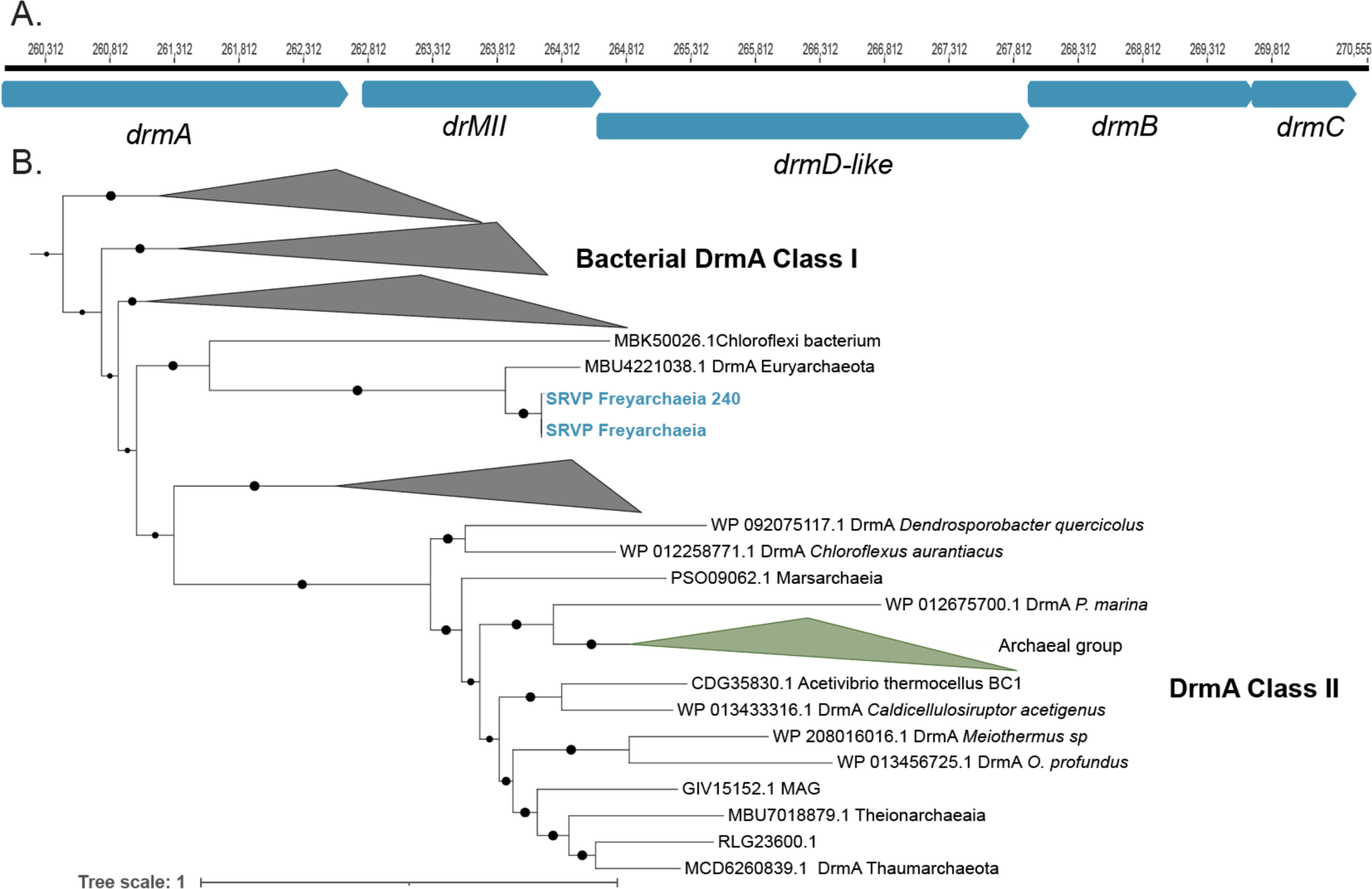
Genomic organization of DISARM (Defense Island System Associated with Restriction-Modification) system from Freyarchaeia and phylogenetic placement of the A DrmA within the whole superfamily. **A**. DrmA, DrmMII, DrmA’, DrmB and DrmC gene locus. **B.** Phylogenetic analysis of the DrmA protein found within the Freyarchaeia genome and reference sequences. The phylogenetic tree was initially constructed with IQ-TREE (version 1.6.1) employing the LG+FO+R model.

**Figure S8.**
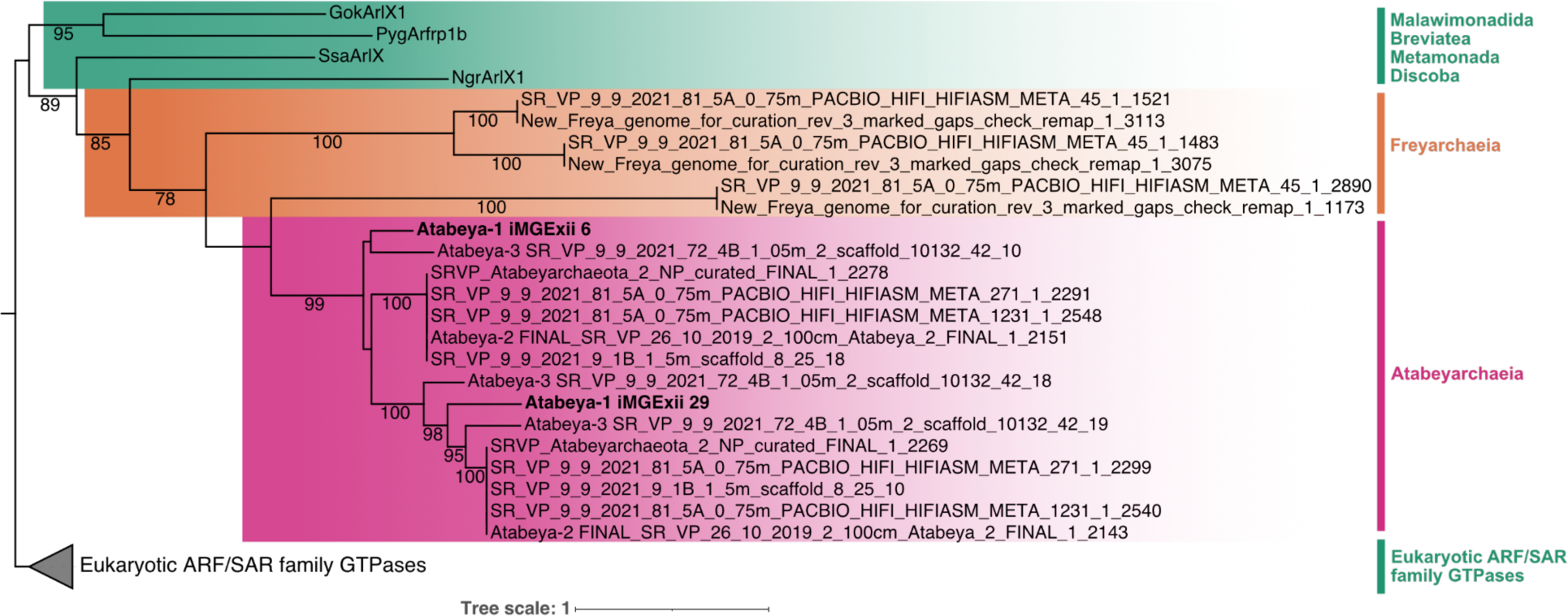
Comparison of GTPases encoded within Atabeyarchaeia putative MGEs and those integrated into the genomes of other Asgard archaea and Eukaryotes. Maximum likelihood tree of eukaryotic ARF family GTPases and Asgard archaea GTPase sequences. GTPases on the putative iMGE-xii are highlighted in bold. A curated set of proteins for the eukaryotic Arf family described in Vargová et al., (2021) was used to search in both Atabeyarchaeia and Freyarchaeia genomes and MGEs. A non-redundant subset of the references and Asgard hits with greater than 25% protein identity were aligned and trimmed with Mafft auto (v7.310) and trimAl-gt 0.5 (v1.4.rev15). The final tree was produced with iqtree (v.1.6.1) and LG+R9 model was chosen according to BIC.

**Figure S9.**
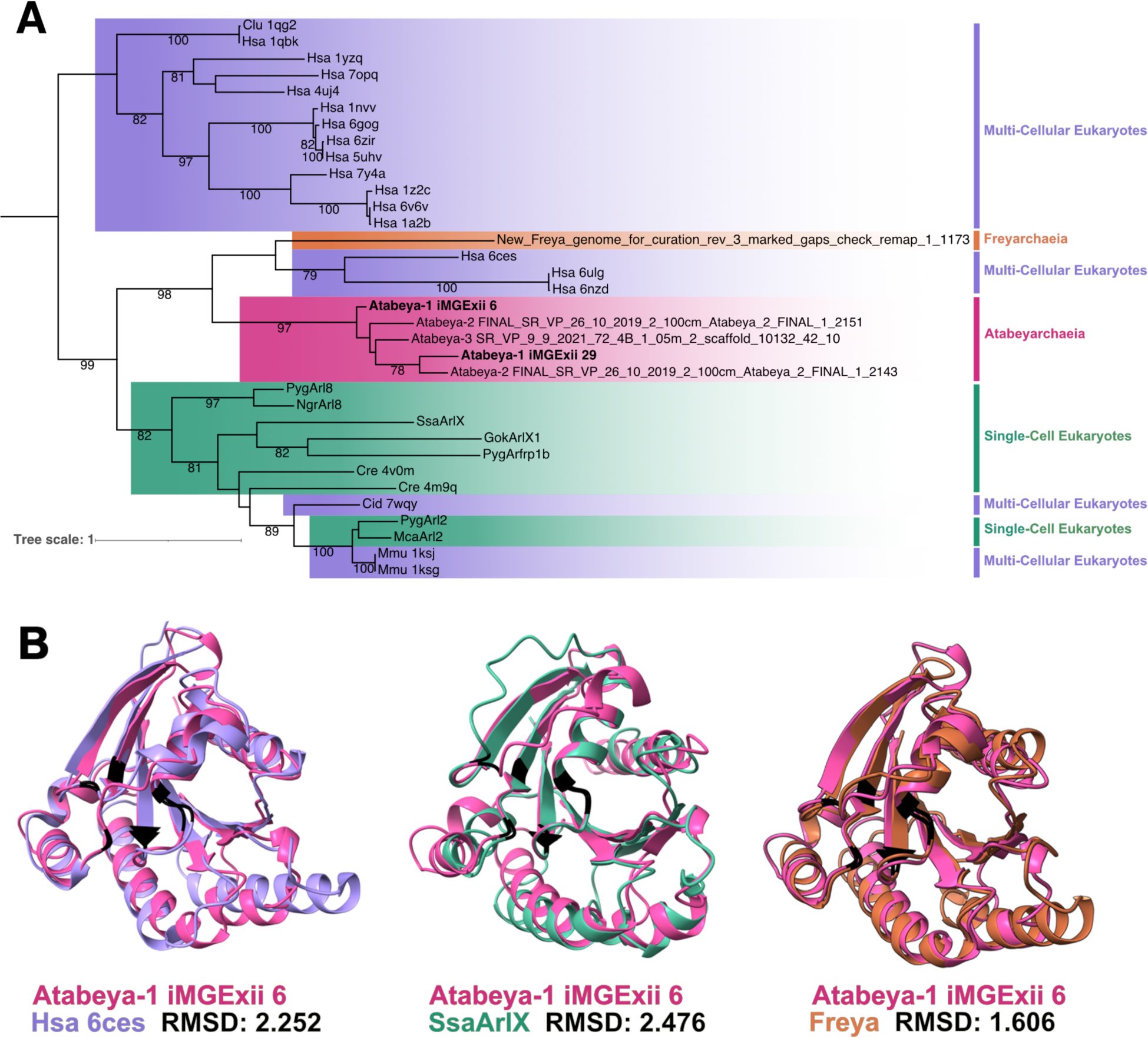
Structural similarity between Atabeyarchaeia MGE and eukaryotic GTPases. **(A)** Structural phylogeny including a subset of the proteins with the sequence phylogeny and confirmed structural homologues in the RCSB Protein Data Bank (See “Methods”). The first three letters of the references in sequence and structural phylogenies are the first letter of the genus followed by the first two letters of the species (i.e., Hsa abbreviated for *Homo sapien*). **(B)** Atabeya-1 iMGE-xii_6 GTPase structural model (pink) superimposed on a *H. sapien* GTPase predicted by electron microscopy (purple), *Spironucleus salmonicida* GTPase structural model (green), and Freyarchaeia GTPase structural model (orange) (from left to right).

**Figure S10.**
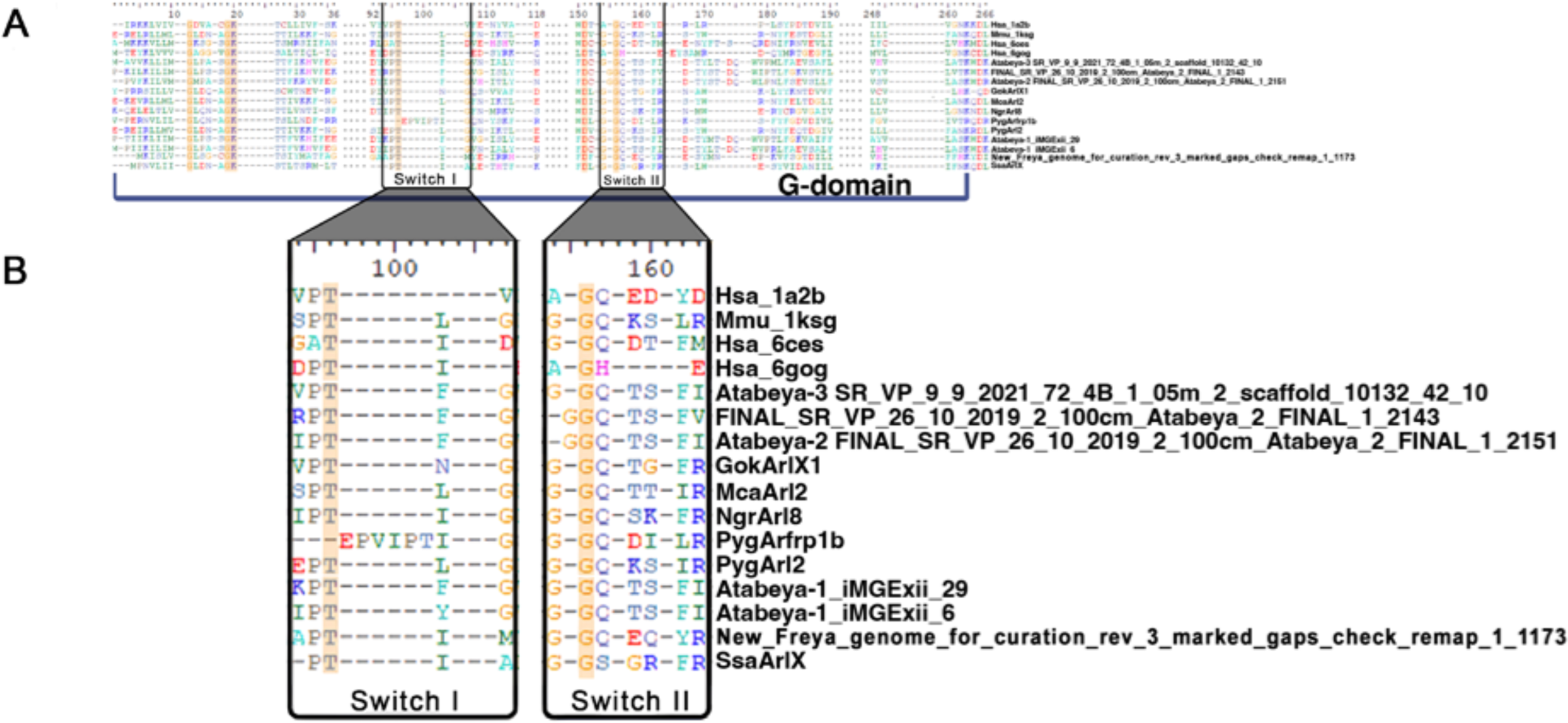
Conservation of G domain and switches. **(A)** Multi-sequence structural alignment, highlighting the conserved amino acids (yellow) in the G domain and **(B)** the position of the switches.

## References

Al-Shayeb, B., Schoelmerich, M. C., West-Roberts, J., Valentin-Alvarado, L. E., Sachdeva, R., Mullen, S., Crits-Christoph, A., Wilkins, M. J., Williams, K. H., Doudna, J. A., & Banfield, J. F. (2022). Borgs are giant genetic elements with potential to expand metabolic capacity. Nature, 610(7933), 731–736.

Anton, B. P., & Roberts, R. J. (2021). Beyond Restriction Modification: Epigenomic Roles of DNA Methylation in Prokaryotes. Annual Review of Microbiology, 75, 129–149.

Berman, H. M., Westbrook, J., Feng, Z., Gilliland, G., Bhat, T. N., Weissig, H., Shindyalov, I. N., & Bourne, P. E. (2000). The Protein Data Bank. Nucleic Acids Research, 28(1), 235– 242.

Blow, M. J., Clark, T. A., Daum, C. G., Deutschbauer, A. M., Fomenkov, A., Fries, R., Froula, J., Kang, D. D., Malmstrom, R. R., Morgan, R. D., Posfai, J., Singh, K., Visel, A., Wetmore, K., Zhao, Z., Rubin, E. M., Korlach, J., Pennacchio, L. A., & Roberts, R. J. (2016). The Epigenomic Landscape of Prokaryotes. PLoS Genetics, 12(2), e1005854.

Bravo, J. P. K., Aparicio-Maldonado, C., Nobrega, F. L., Brouns, S. J. J., & Taylor, D. W. (2022). Structural basis for broad anti-phage immunity by DISARM. Nature Communications, 13(1), 2987.

Bushnell, B. (2014). BBMap: A Fast, Accurate, Splice-Aware Aligner (No. LBNL-7065E). Lawrence Berkeley National Lab. (LBNL), Berkeley, CA (United States). https://www.osti.gov/servlets/purl/1241166

Chen, L.-X., Anantharaman, K., Shaiber, A., Eren, A. M., & Banfield, J. F. (2020). Accurate and complete genomes from metagenomes. Genome Research, 30(3), 315–333.

Dong, R., Peng, Z., Zhang, Y., & Yang, J. (2018). mTM-align: an algorithm for fast and accurate multiple protein structure alignment. Bioinformatics, 34(10), 1719–1725.

Doron, S., Melamed, S., Ofir, G., Leavitt, A., Lopatina, A., Keren, M., Amitai, G., & Sorek, R. (2018). Systematic discovery of antiphage defense systems in the microbial pangenome. Science, 359(6379). 10.1126/science.aar4120

Eme, L., Tamarit, D., Caceres, E. F., Stairs, C. W., De Anda, V., Schön, M. E., Seitz, K. W., Dombrowski, N., Lewis, W. H., Homa, F., Saw, J. H., Lombard, J., Nunoura, T., Li, W.-J., Hua, Z.-S., Chen, L.-X., Banfield, J. F., John, E. S., Reysenbach, A.-L., … Ettema, T. J. G. (2023). Inference and reconstruction of the heimdallarchaeial ancestry of eukaryotes. Nature, 618(7967), 992–999.

Feng, X., Cheng, H., Portik, D., & Li, H. (2022). Metagenome assembly of high-fidelity long reads with hifiasm-meta. Nature Methods, 19(6), 671–674.

Ghaly, T. M., Tetu, S. G., Penesyan, A., Qi, Q., Rajabal, V., & Gillings, M. R. (2022). Discovery of integrons in Archaea: Platforms for cross-domain gene transfer. Science Advances, 8(46), eabq6376.

Gomis-Rüth, F. X., Solá, M., Acebo, P., Párraga, A., Guasch, A., Eritja, R., González, A., Espinosa, M., del Solar, G., & Coll, M. (1998). The structure of plasmid-encoded transcriptional repressor CopG unliganded and bound to its operator. The EMBO Journal, 17(24), 7404–7415.

Guo, X., & Huang, L. (2010). A superfamily 3 DNA helicase encoded by plasmid pSSVi from the hyperthermophilic archaeon Sulfolobus solfataricus unwinds DNA as a higher-order oligomer and interacts with host primase. Journal of Bacteriology, 192(7), 1853–1864.

Hug, L. A., Castelle, C. J., Wrighton, K. C., Thomas, B. C., Sharon, I., Frischkorn, K. R., Williams, K. H., Tringe, S. G., & Banfield, J. F. (2013). Community genomic analyses constrain the distribution of metabolic traits across the Chloroflexi phylum and indicate roles in sediment carbon cycling. Microbiome, 1(1), 22.

Jumper, J., Evans, R., Pritzel, A., Green, T., Figurnov, M., Ronneberger, O., Tunyasuvunakool, K., Bates, R., Žídek, A., Potapenko, A., Bridgland, A., Meyer, C., Kohl, S. A. A., Ballard, A. J., Cowie, A., Romera-Paredes, B., Nikolov, S., Jain, R., Adler, J., … Hassabis, D. (2021). Highly accurate protein structure prediction with AlphaFold. Nature, 596(7873), 583–589.

Kieft, K., & Anantharaman, K. (2022). Deciphering Active Prophages from Metagenomes. mSystems, 7(2), e0008422.

Krupovic, M., Makarova, K. S., Wolf, Y. I., Medvedeva, S., Prangishvili, D., Forterre, P., & Koonin, E. V. (2019). Integrated mobile genetic elements in Thaumarchaeota. Environmental Microbiology, 21(6), 2056–2078.

Langmead, B., Trapnell, C., Pop, M., & Salzberg, S. L. (2009). Ultrafast and memory-efficient alignment of short DNA sequences to the human genome. Genome Biology, 10(3), R25.

Medvedeva, S., Sun, J., Yutin, N., Koonin, E. V., Nunoura, T., Rinke, C., & Krupovic, M. (2022). Three families of Asgard archaeal viruses identified in metagenome-assembled genomes. Nature Microbiology, 7(7), 962–973.

Meng, E. C., Goddard, T. D., Pettersen, E. F., Couch, G. S., Pearson, Z. J., Morris, J. H., & Ferrin, T. E. (2023). UCSF ChimeraX: Tools for structure building and analysis. Protein Science: A Publication of the Protein Society, 32(11), e4792.

Mirdita, M., Schütze, K., Moriwaki, Y., Heo, L., Ovchinnikov, S., & Steinegger, M. (2022). ColabFold: making protein folding accessible to all. Nature Methods, 19(6), 679–682.

Mizuno, C. M., Prajapati, B., Lucas-Staat, S., Sime-Ngando, T., Forterre, P., Bamford, D. H., Prangishvili, D., Krupovic, M., & Oksanen, H. M. (2019). Novel haloarchaeal viruses from Lake Retba infecting Haloferax and Halorubrum species. Environmental Microbiology, 21(6), 2129–2147.

Moreno-Cinos, C., Goossens, K., Salado, I. G., Van Der Veken, P., De Winter, H., & Augustyns, K. (2019). ClpP Protease, a Promising Antimicrobial Target. International Journal of Molecular Sciences, 20(9). 10.3390/ijms20092232

Morgan, R. D., Dwinell, E. A., Bhatia, T. K., Lang, E. M., & Luyten, Y. A. (2009). The MmeI family: type II restriction-modification enzymes that employ single-strand modification for host protection. Nucleic Acids Research, 37(15), 5208–5221.

Ofir, G., Melamed, S., Sberro, H., Mukamel, Z., Silverman, S., Yaakov, G., Doron, S., & Sorek, R. (2018). DISARM is a widespread bacterial defence system with broad anti-phage activities. Nature Microbiology, 3(1), 90–98.

Payne, L. J., Meaden, S., Mestre, M. R., Palmer, C., Toro, N., Fineran, P. C., & Jackson, S. A. (2022). PADLOC: a web server for the identification of antiviral defence systems in microbial genomes. Nucleic Acids Research, 50(W1), W541–W550.

Rambo, I. M., Langwig, M. V., Leão, P., De Anda, V., & Baker, B. J. (2022). Genomes of six viruses that infect Asgard archaea from deep-sea sediments. Nature Microbiology, 7(7), 953–961.

Raymann, K., Forterre, P., Brochier-Armanet, C., & Gribaldo, S. (2014). Global phylogenomic analysis disentangles the complex evolutionary history of DNA replication in archaea. Genome Biology and Evolution, 6(1), 192–212.

Roberts, R. J., Vincze, T., Posfai, J., & Macelis, D. (2015). REBASE--a database for DNA restriction and modification: enzymes, genes and genomes. Nucleic Acids Research, 43(Database issue), D298–D299.

Rocha, E. P. C., & Bikard, D. (2022). Microbial defenses against mobile genetic elements and viruses: Who defends whom from what? PLoS Biology, 20(1), e3001514.

Rubin, B. E., Diamond, S., Cress, B. F., Crits-Christoph, A., Lou, Y. C., Borges, A. L., Shivram, H., He, C., Xu, M., Zhou, Z., Smith, S. J., Rovinsky, R., Smock, D. C. J., Tang, K., Owens, T. K., Krishnappa, N., Sachdeva, R., Barrangou, R., Deutschbauer, A. M., … Doudna, J. A. (2022). Species- and site-specific genome editing in complex bacterial communities. Nature Microbiology, 7(1), 34–47.

Russel, J., Pinilla-Redondo, R., Mayo-Muñoz, D., Shah, S. A., & Sørensen, S. J. (2020). CRISPRCasTyper: Automated Identification, Annotation, and Classification of CRISPR-Cas Loci. The CRISPR Journal, 3(6), 462–469.

Seitz, K. W., Dombrowski, N., Eme, L., Spang, A., Lombard, J., Sieber, J. R., Teske, A. P., Ettema, T. J. G., & Baker, B. J. (2019). Asgard archaea capable of anaerobic hydrocarbon cycling. Nature Communications, 10(1), 1822.

Senčilo, A., & Roine, E. (2014). A Glimpse of the genomic diversity of haloarchaeal tailed viruses. Frontiers in Microbiology, 5, 84.

Shen, B. W., Heiter, D. F., Chan, S.-H., Wang, H., Xu, S.-Y., Morgan, R. D., Wilson, G. G., & Stoddard, B. L. (2010). Unusual target site disruption by the rare-cutting HNH restriction endonuclease PacI. Structure, 18(6), 734–743.

Spang, A., Saw, J. H., Jørgensen, S. L., Zaremba-Niedzwiedzka, K., Martijn, J., Lind, A. E., van Eijk, R., Schleper, C., Guy, L., & Ettema, T. J. G. (2015). Complex archaea that bridge the gap between prokaryotes and eukaryotes. Nature, 521(7551), 173–179.

Speth, D. R., Yu, F. B., Connon, S. A., Lim, S., Magyar, J. S., Peña-Salinas, M. E., Quake, S. R., & Orphan, V. J. (2022). Microbial communities of Auka hydrothermal sediments shed light on vent biogeography and the evolutionary history of thermophily. The ISME Journal, 16(7), 1750–1764.

Takai, Y., Sasaki, T., & Matozaki, T. (2001). Small GTP-binding proteins. Physiological Reviews, 81(1), 153–208.

Tamarit, D., Caceres, E. F., Krupovic, M., Nijland, R., Eme, L., Robinson, N. P., & Ettema, T. J. G. (2022). A closed Candidatus Odinarchaeum chromosome exposes Asgard archaeal viruses. Nature Microbiology, 7(7), 948–952.

Tesson, F., Hervé, A., Mordret, E., Touchon, M., d’Humières, C., Cury, J., & Bernheim, A. (2022). Systematic and quantitative view of the antiviral arsenal of prokaryotes. Nature Communications, 13(1), 2561.

Valentin-Alvarado, L. E., Appler, K. E., De Anda, V., Schoelmerich, M. C., West-Roberts, J., Kivenson, V., Crits-Christoph, A., Ly, L., Sachdeva, R., Savage, D. F., Baker, B. J., & Banfield, J. F. (2023). Asgard archaea modulate potential methanogenesis substrates in wetland soil. In bioRxiv (p. 2023.11.21.568159). 10.1101/2023.11.21.568159

van Kempen, M., Kim, S. S., Tumescheit, C., Mirdita, M., Lee, J., Gilchrist, C. L. M., Söding, J., & Steinegger, M. (2023). Fast and accurate protein structure search with Foldseek. Nature Biotechnology. 10.1038/s41587-023-01773-0

Vetter, I. R., & Wittinghofer, A. (2001). The guanine nucleotide-binding switch in three dimensions. Science, 294(5545), 1299–1304.

Wein, T., & Sorek, R. (2022). Bacterial origins of human cell-autonomous innate immune mechanisms. Nature Reviews. Immunology, 22(10), 629–638.

Wu, F., Speth, D. R., Philosof, A., Crémière, A., Narayanan, A., Barco, R. A., Connon, S. A., Amend, J. P., Antoshechkin, I. A., & Orphan, V. J. (2022). Unique mobile elements and scalable gene flow at the prokaryote-eukaryote boundary revealed by circularized Asgard archaea genomes. Nature Microbiology, 7(2), 200–212.

Zaremba-Niedzwiedzka, K., Caceres, E. F., Saw, J. H., Bäckström, D., Juzokaite, L., Vancaester, E., Seitz, K. W., Anantharaman, K., Starnawski, P., Kjeldsen, K. U., Stott, M. B., Nunoura, T., Banfield, J. F., Schramm, A., Baker, B. J., Spang, A., & Ettema, T. J. G. (2017). Asgard archaea illuminate the origin of eukaryotic cellular complexity. Nature, 541(7637), 353– 358.

