## Supplementary Information for "Genetic elements and defense systems drive diversification and evolution in Asgard archaea"

| Mobile Genetic Element | MGE type | Host | Genome size (Kb) |
|---|---|---|---|
| iMGE-i Yucahu | Plasmid/integrase | Ata-1 | 10.9 |
| iMGE-i mini-Yucahu | unknown | Ata-1 | 2.6 |
| iMGE-ii | Putative transposase | Ata-1 | 6.3 |
| iMGE-iii | Putative transposase | Ata-1 | 4.4 |
| iMGE-iv | Transposase-IS605 | Ata-1 | 5.3 |
| iMGE-v | viral/mobile/integrase | Ata-1 | 26.8 |
| iMGE-vi | viral/mobile/integrase | Ata-1 | 22.9 |
| iMGE-vii | viral/mobile/integrase | Ata-1 | 12.9 |
| iMGE-viii | Transposase-IS605 | Ata-1 | 1.3 |
| iMGE-ix | Transposase-IS605 | Ata-1 | 9.3 |
| iMGE-x | viral/mobile | Ata-1 | 12 |
| iMGE-xi | viral/mobile/integrase | Ata-1 | 20.5 |
| iMGE-xii | unknown | Ata-1 | 25.8 |
| iMGE-xiii | mobilome /transposase | Ata-1 | 2.5 |
| iMGE-xiv | mobilome /transposase | Ata-1 | 1.0 |
| iMGE-xv | unknown | Ata-2 | 18 |
| iMGE-xvi | Plasmid-like | Ata-2 | 8.0 |
| iMGE-xvii | mobile/integrase | Ata-2 | 9.5 |
| iMGE-xviii | viral/mobile | Ata-2 | 29.5 |
| Opia-2829 (reference genome) | Tailed virus | Ata-2 | 40.0 |
| Opia-3708 related genotype | Tailed virus | Ata-2 | 32.8 |
| MGE-1318 | novel MGE | Ata-1 | 26.3 |
| MGE-9917 (related to MGE-1318) | novel MGE | Ata-1 | 26.7 |
| Plasmid-700 | Plasmid | Ata-2 | 28.8 |

**Table S1: Manually curated genomes for chromosomally integrated genetic elements and extrachromosomal elements**. MGE type is the closest classification based on gene content and genome structure. The host was determined based on matches of the MGE squences to spacers from the CRISPR systems.

| Sample | motifString | modificationType |
|---|---|---|
| Atabeyarchaeia 1 | GNNGANNNNNNNRTTC | m6A |
| Atabeyarchaeia 1 | CCGG | m4C |
| Atabeyarchaeia 1 | GGCC | m4C |
| Atabeyarchaeia 1 | CGCG | m4C |
| Atabeyarchaeia 1 | GTSAC | m6A |
| Atabeyarchaeia 1 | CTAG | m4C |
| Atabeyarchaeia 1 | CTSAG | m6A |
| Atabeyarchaeia 1 | CCAGG | m6A |
| Atabeyarchaeia 1 | DGCGCH | m4C |
| Atabeyarchaeia 1 | GAGGAA | m6A |
| Atabeyarchaeia 1 | GATC | m6A |
| Atabeyarchaeia 1 | GTAC | m4C |
| Atabeyarchaeia 1 | GYATGAG | m6A |
| Atabeyarchaeia 2 | GTSAC | m6A |
| Atabeyarchaeia 2 | CAGNNNNNRTGG | m6A |
| Atabeyarchaeia 2 | CGCG | m4C |
| Atabeyarchaeia 2 | GATC | m6A |
| Atabeyarchaeia 2 | CTAG | m4C |
| Atabeyarchaeia 2 | ACAGG | m6A |
| Atabeyarchaeia 2 | AGSCT | m4C |
| Atabeyarchaeia 2 | DGCGCH | m4C |
| Atabeyarchaeia 2 | GGCAG | m6A |
| Atabeyarchaeia 2 | GRATGAG | m6A |
| Atabeyarchaeia 2 | GTAC | m4C |
| Freyarchaeia strain | CCGG | m4C |
| Freyarchaeia strain | GGCC | m4C |
| Freyarchaeia strain | CCWGG | m4C |
| Freyarchaeia strain | GCGC | m4C |
| Freyarchaeia strain | GGWCC | m4C |
| Freyarchaeia strain | CCGG | m4C |

**Table S2** Methylation patterns in Atabeyarchaeia and Freyarchaeia genomes based on PacBio data.

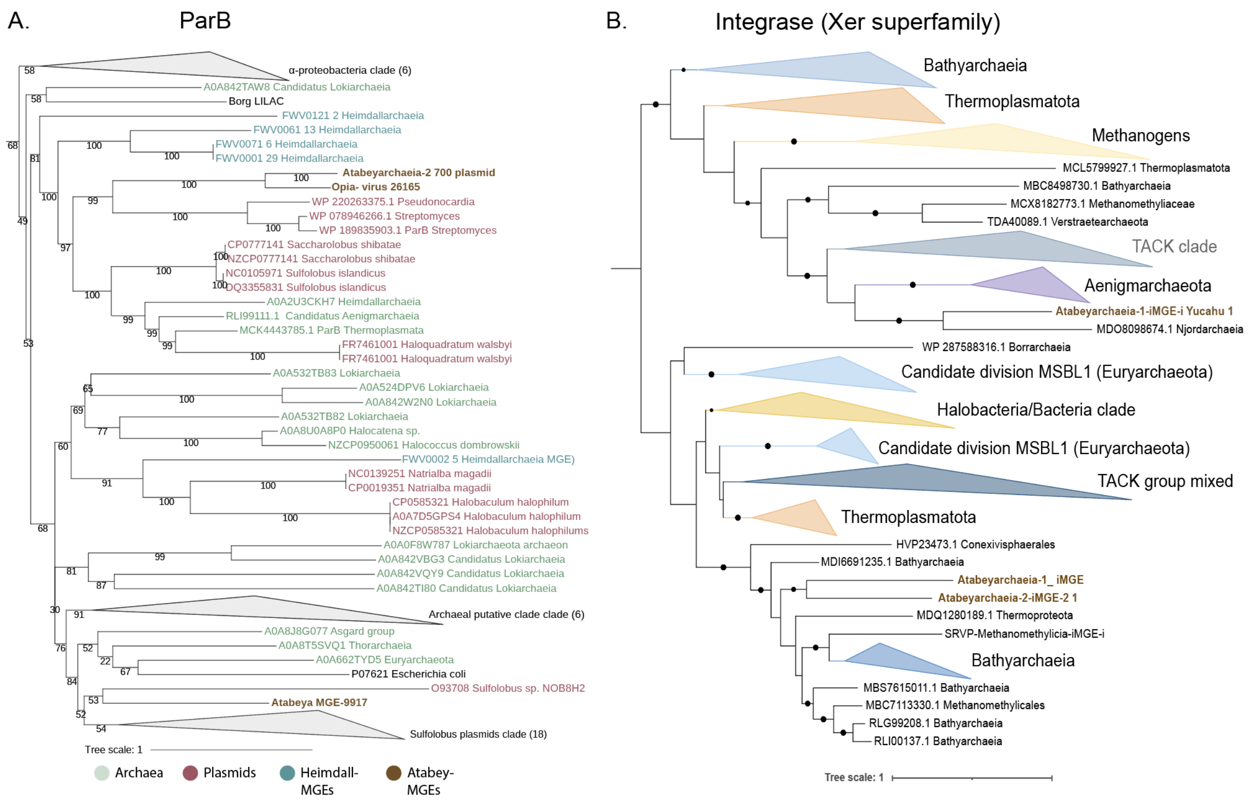

**Figure S1: Phylogenetic analyses of ParB and tyrosine-recombinase-like proteins across archaeal mobile genetic elements. A.** Phylogenetic tree for ParB and ParB-like Proteins. Analysis incorporates reference sequences of *E. coli* ParB (Pfam08775) and *Sulfolobus* conjugative plasmid ParB (NOB8H2), alongside representative ParB-like domains from select archaeal and bacterial genomes. Selection criteria for inclusion involved a rigorous screening of the top 25 protein hits from the NCBI database. Sequence alignments were conducted using MAFFT (v7.310) in 'auto' mode, with subsequent optimization by trimAl (v1.4.rev15) applying a 0.7 gap threshold. Phylogenetic trees were initially inferred using IQ-TREE (v1.6.1) under the LG+FO+R model. This segment details **B.** Phylogeny of tyrosine-recombinase-like proteins within integrated genetic elements of Atabeyarchaeia-1 and Atabeyarchaeia-2. The analysis selected the top 25 protein hits from the NCBI database, aligned with MAFFT (version 7.310) on 'auto' setting, and refined the alignment with trimAl (version 1.4.rev15) using a 0.7 gap threshold. The phylogenetic framework was constructed using IQ-TREE (version 1.6.1), adopting the LG+FO+R model.

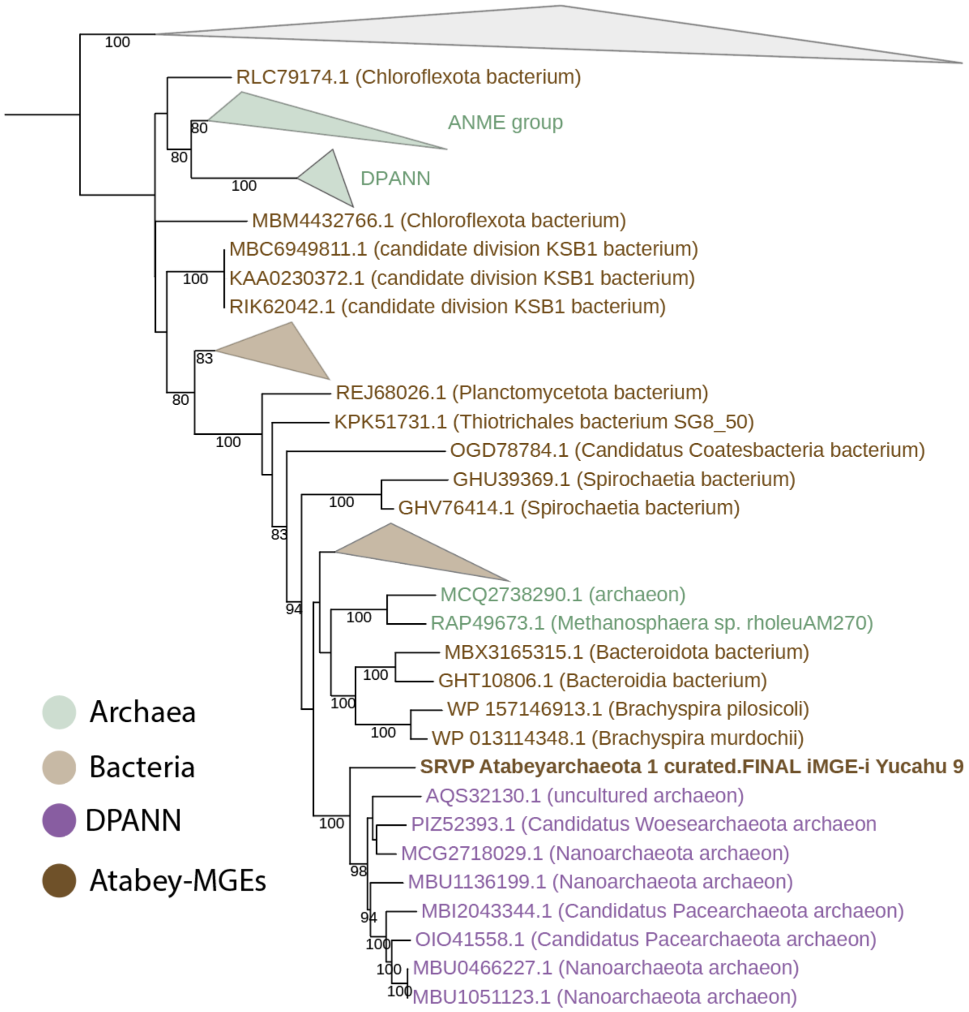

**Figure S2: Maximum likelihood phylogeny of Type II-G restriction-modification (IIG RM) protein fusion that combines endonuclease and methyltransferase present in Yucahy MGE** The phylogenetic tree was initially constructed with IQ-TREE (version 1.6.1) employing the LG+FO+R model for the first iterations. Subsequent refinement involved several rounds of manual branch checking to ensure the accuracy and reliability of the phylogenetic relationships depicted.

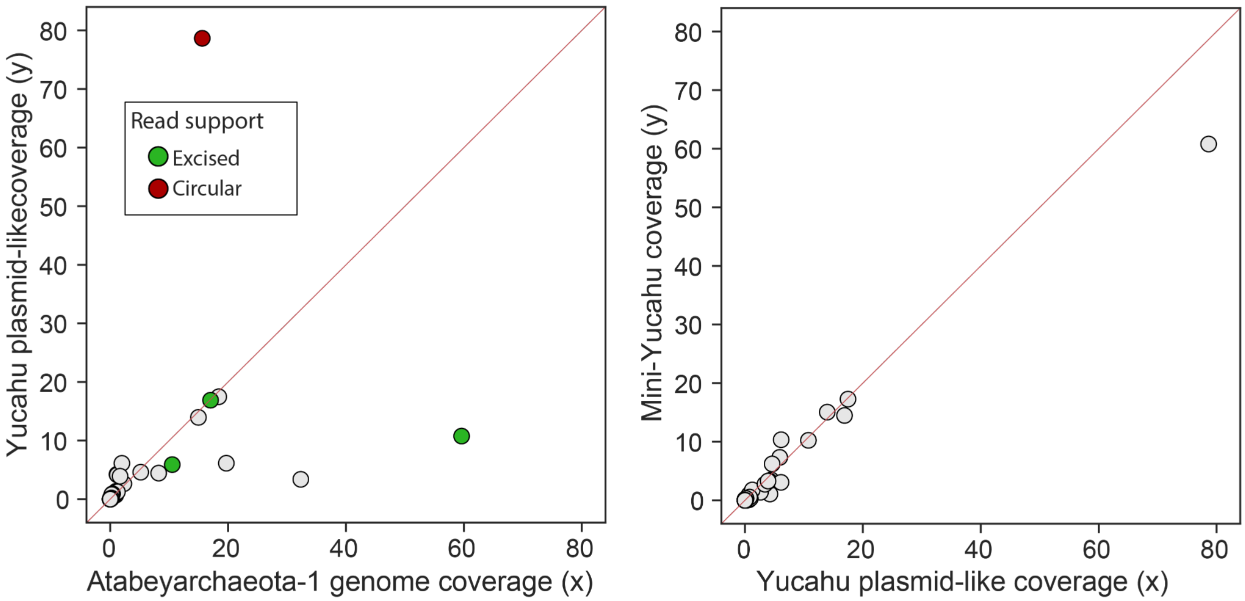

**Figure S3: Cross-sample Atabeyarchaeia-1 plasmid integration A.** Correlations of the genome coverages between integrated plasmid Yucahu and Atabeyarchaeia **B.** Correlations of the Yucahu plasmid genome and the mini-Yucahu element

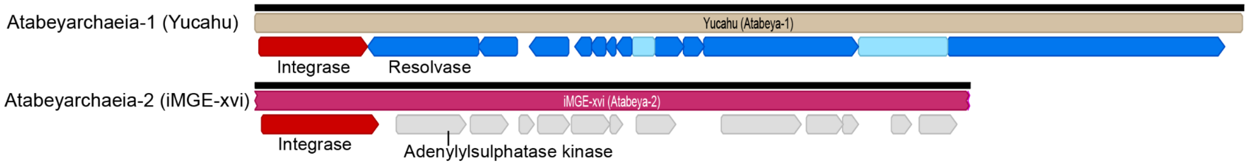

**Figure S4 Comparison of gene content of Yucahu-i (Atabeyarchaeia-1), top. and iMGE**-xvi **(Atabeyarchaeia-2), bottom.** Annotated proteins of Yucahu-i include a tyrosine recombinase and a protein with similarity to adenylylsulfate kinases. The iMGE-xvi has 12 open reading frames, most encoding hypothetical proteins or proteins of unknown function. Three proteins are predicted to have 5 - 8 transmembrane domains. One is a putative membrane-bound serine protease of the ClpP class involved in the proteolysis of misfolded and defective proteins(Moreno-Cinos et al., 2019)**.**

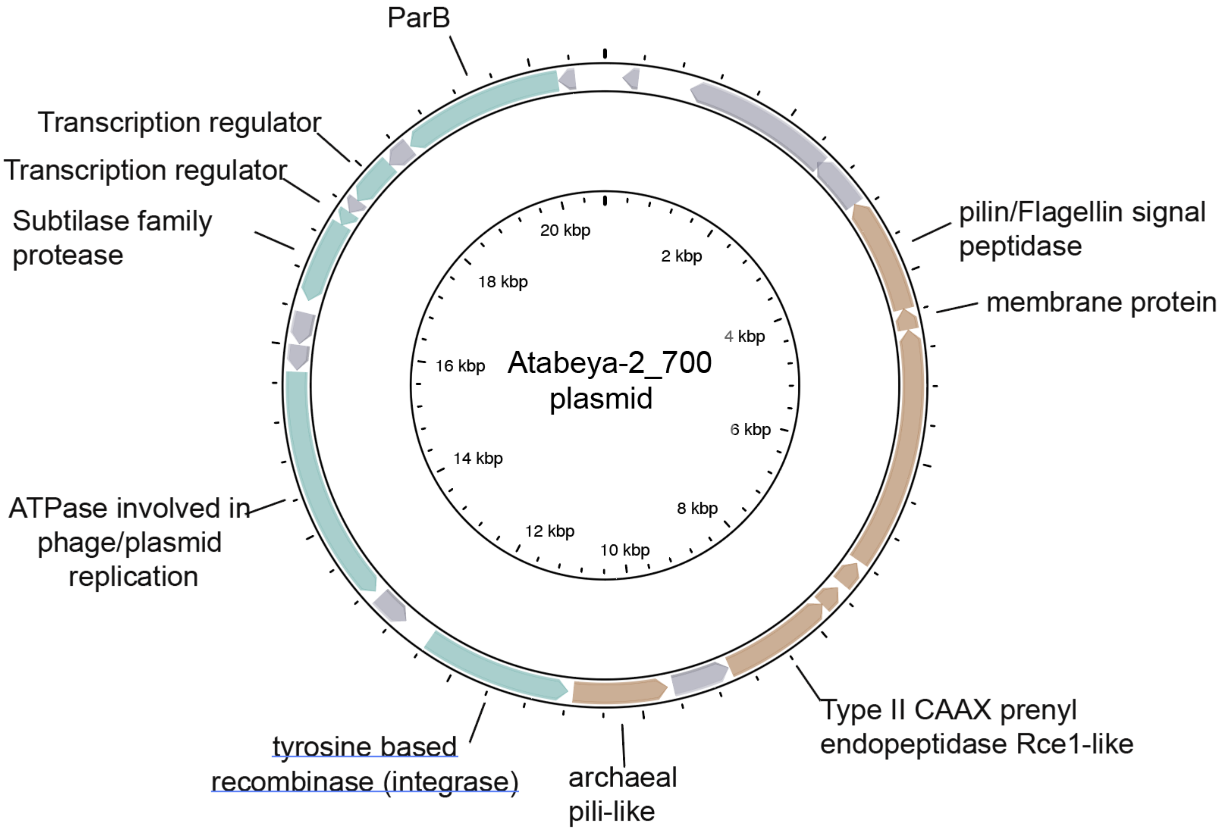

**Figure S5. Genome diagram of Atabeyarchaeia-2 putative plasmid scaffold_700.** The plasmid has 23 open reading frames and gray-colored genes are hypothetical. Genes colored brown have transmembrane domains and are predicted to be extracellular.

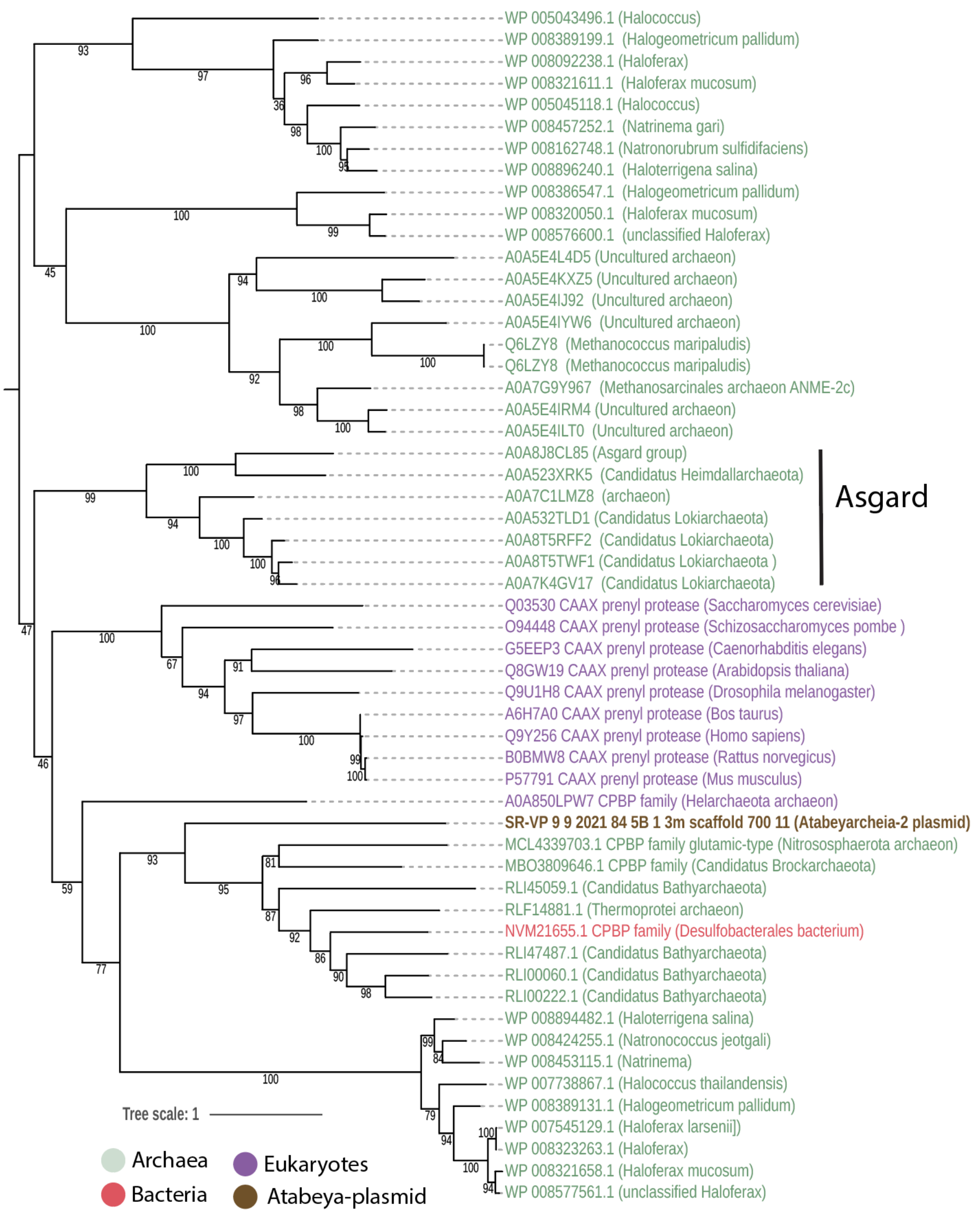

**Figure S6: Maximum likelihood phylogeny of CAAX proteases identified in the Atabeyarchaeia-2 plasmid_700.** For this analysis, the top 50 protein hits from the NCBI database were selected. Sequence alignment was performed using MAFFT (version 7.310) with the 'auto' setting, and the alignment was subsequently trimmed with trimAl (version 1.4.rev15) using a gap threshold of 0.7 (-gt 0.7). The phylogenetic tree was initially constructed with IQ-TREE (version 1.6.1) employing the LG+FO+R model.

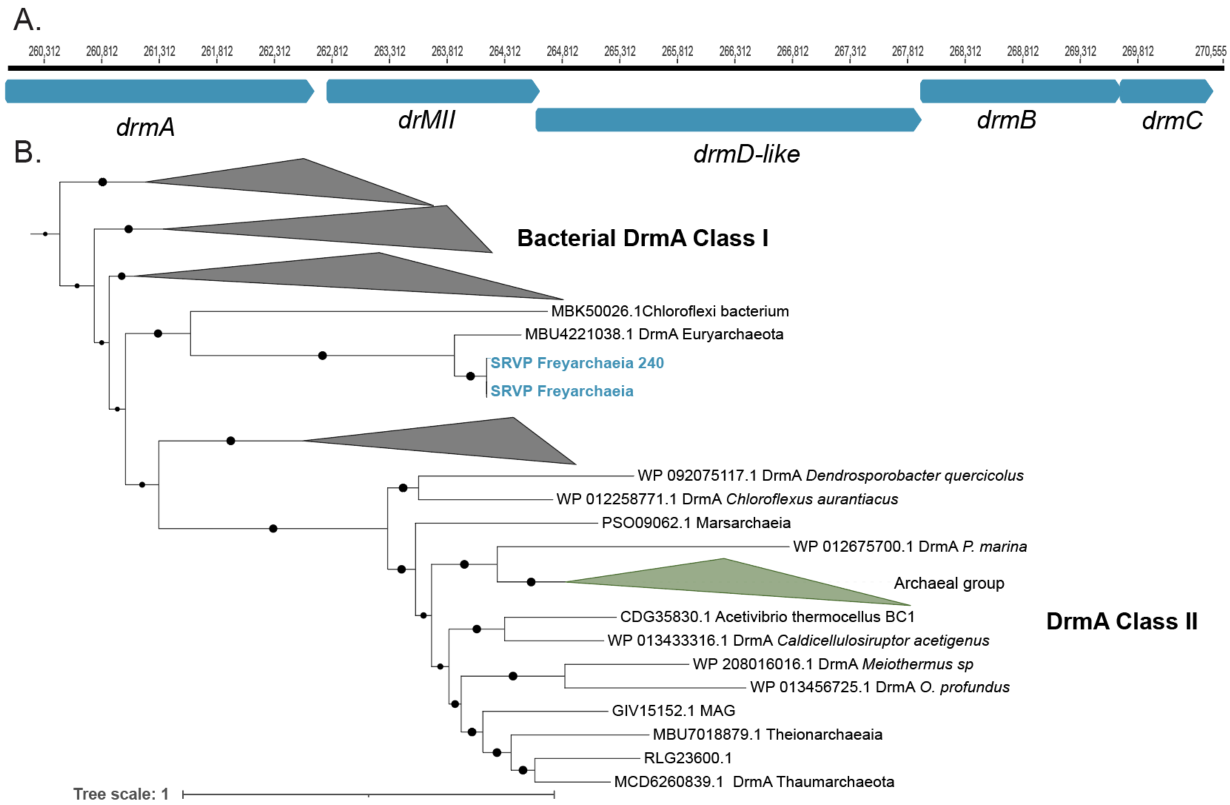

**Figure S7: Genomic organization of DISARM (Defense Island System Associated with Restriction-Modification) system from Freyarchaeia and phylogenetic placement of the A DrmA within the whole superfamily. A.** DrmA, DrmMII, DrmA', DrmB and DrmC gene locus. **B.** Phylogenetic analysis of the DrmA protein found within the Freyarchaeia genome and reference sequences. The phylogenetic tree was initially constructed with IQ-TREE (version 1.6.1) employing the LG+FO+R model.

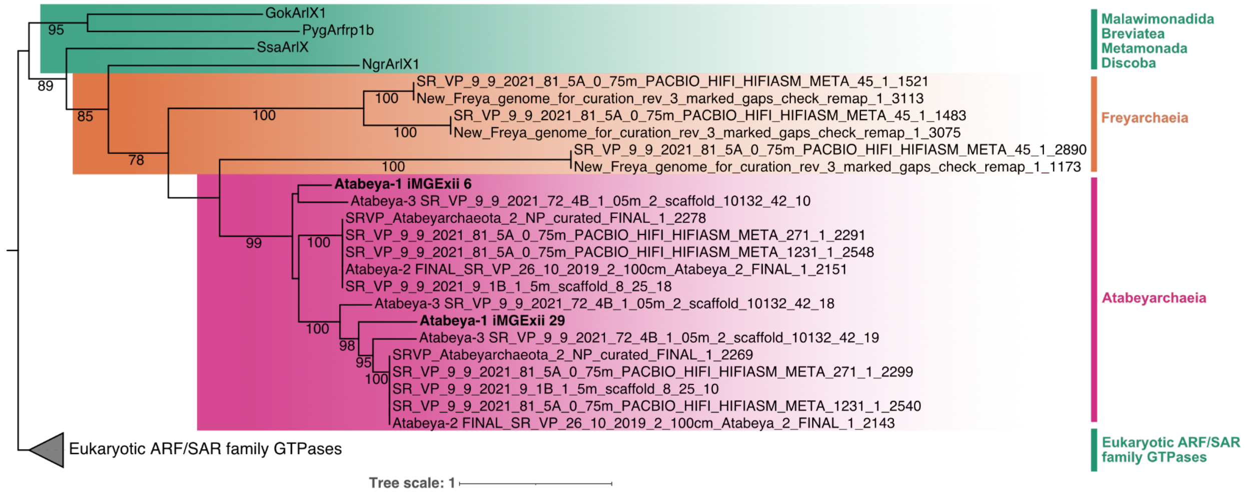

**Figure S8 Comparison of GTPases encoded within Atabeyarchaeia putative MGEs and those integrated into the genomes of other Asgard archaea and Eukaryotes.** Maximum likelihood tree of eukaryotic ARF family GTPases and Asgard archaea GTPase sequences. GTPases on the putative iMGE-xii are highlighted in bold. A curated set of proteins for the eukaryotic Arf family described in Vargová et al., (2021) was used to search in both Atabeyarchaeia and Freyarchaeia genomes and MGEs. A non-redundant subset of the references and Asgard hits with greater than 25% protein identity were aligned and trimmed with Mafft auto (v7.310) and trimAl-gt 0.5 (v1.4.rev15). The final tree was produced with iqtree (v.1.6.1) and LG+R9 model was chosen according to BIC.

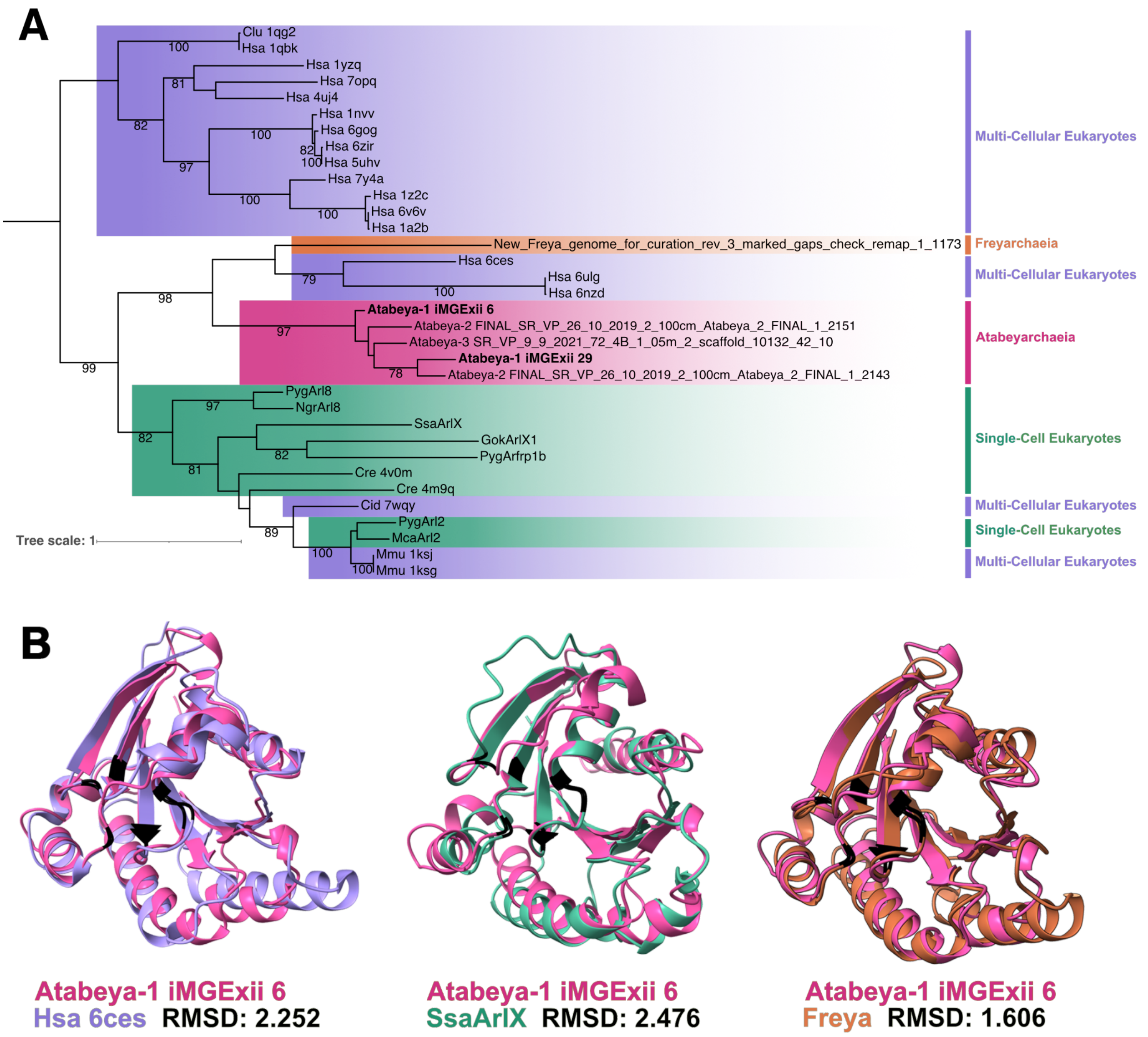

**Figure S9 Structural similarity between Atabeyarchaeia MGE and eukaryotic GTPases. (A)** Structural phylogeny including a subset of the proteins with the sequence phylogeny and confirmed structural homologues in the RCSB Protein Data Bank (See "Methods"). The first three letters of the references in sequence and structural phylogenies are the first letter of the genus followed by the first two letters of the species (i.e., Hsa abbreviated for *Homo sapien*). **(B)** Atabeya-1 iMGE-xii_6 GTPase structural model (pink) superimposed on a *H. sapien* GTPase predicted by electron microscopy (purple), *Spironucleus salmonicida* GTPase structural model (green), and Freyarchaeia GTPase structural model (orange) (from left to right).

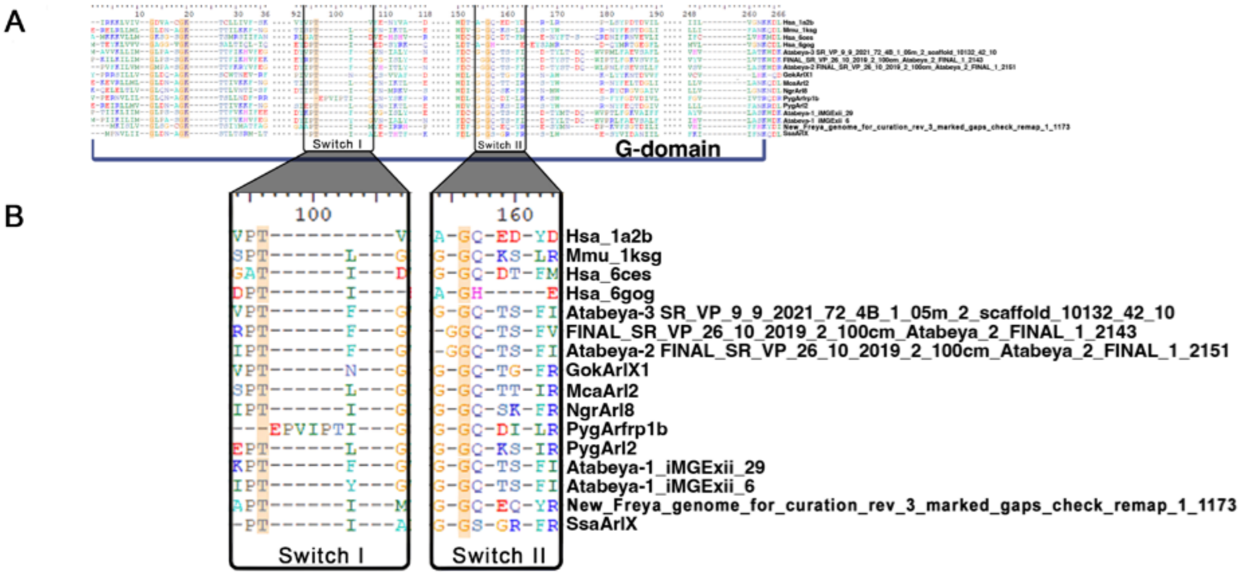

**Figure S10 Conservation of G domain and switches. (A)** Multi-sequence structural alignment, highlighting the conserved amino acids (yellow) in the G domain and **(B)** the position of the switches.
